# Attention modulates neural representations of acoustic, categorical, and identity features in a task-dependent manner

**DOI:** 10.64898/2026.01.07.698102

**Authors:** Onnipekka V. Varis, Ilkka A. Muukkonen, Patrik A. Wikman

**Affiliations:** Department of Psychology, PO Box 21 FI-00014 University of Helsinki, Finland; Neuroscience Center, HiLife, PO Box 21 FI-00014 University of Helsinki, Finland; Department of Brain and Cognition, PO box 03711, KU Leuven, Leuven, Belgium; Advanced Magnetic Imaging Centre, PO Box 11000 FI-00076, Aalto University, Espoo, Finland; Helsinki Collegium For Advanced Studies, PO Box 21 FI-00014 University of Helsinki, Finland

## Abstract

The human auditory system represents sounds at multiple levels, from low-level acoustic features to abstract category- and object-level information. Although selective attention enables listening in complex natural soundscapes, it remains unclear which representational levels are modulated by attention and how this depends on scene structure.

Using fMRI, representational similarity analysis, and cross-experiment decoding, we examined attentional modulation of auditory representations under three conditions 1) listening to isolated sounds, 2) attending to one of three overlapping sounds from different categories or 3) one of three overlapping sounds from the same category.

We show that attentional modulation is task-dependent: when competing sounds could not be distinguished by category, attention enhanced low-level acoustic feature processing, whereas in cross-category scenes it targeted abstract category-level representations. Object-identity representations were modulated by attention across both scene types, with category-specific differences, demonstrating flexible, task-dependent targeting of auditory features.

## Introduction

The human auditory system is thought to integrate low-level sound features (e.g., pitch, amplitude) into abstract sound features, which represent complex and meaningful information, such as the category of a sound or the identity of a speaker^1^. These operations have been suggested to rely on neural processing in the auditory ventral stream where sound features are processed along a gradient from simple to abstract^2–4^. Specifically, the primary auditory cortex (A1) differentiates low-level features such as pitch^5^, while the surrounding belt areas are sensitive to band-pass noise bursts—i.e., sounds that contain a range of frequencies within a defined bandwidth^6^. Following the belt regions, the superior temporal gyrus (STG) shows intermediate representations that bridge low-level and abstract sound features^7^. Further along this pathway, both the superior temporal sulcus (STS), anterior portions of the STG and the inferior frontal gyrus (IFG) exhibit category-selective responses to complex sound classes, particularly vocalizations and speech^8–12^.

While the cortical progression from low-level to abstract sound feature processing is relatively well established, much less is known about how attention enables selective processing of naturalistic sound objects in the presence of other competing sounds. A central question in auditory attention research pertains to which sound features are selectively enhanced by attention, and where in the auditory hierarchy these effects emerge. fMRI studies using simple tones have shown that attention can modulate activity already in primary auditory areas in a frequency-specific manner, suggesting that attention may act as a low-level filter^13–16^. However, it is not yet clear whether such effects generalise to other low-level sound features such as amplitude variability or harmonicity.

A limitation of studies using simple stimuli such as tones is that these stimuli do not possess abstract sound features that are present in natural sounds. Indeed, studies using overlapping natural sounds with rich abstract properties (e.g., overlapping speech) have generally not observed reliable low-level attentional effects in early auditory areas. Instead, attentional modulation tends to emerge at later representational stages, typically in STG, STS, and frontal regions^17–21^. For example, in a two-speaker cocktail-party task, attention selectively enhanced articulatory and semantic, but not spectral, feature representations, primarily in higher-order auditory regions^17^. This preferential enhancement of higher-level articulatory and semantic information over low-level spectral coding suggests that, in complex natural scenes, attention often operates at the level of auditory objects or streams rather than individual low-level features.

An important constraint on the generalizability of previous work on auditory attention is that it has largely focused on scenes containing either overlapping simple sounds or overlapping speech. The studies that have investigated attentional modulation in more variable soundscapes suggest attentional modulation is category-specific and emerges in distinct temporal regions depending on the category of the attended sound^20,24,25^. Moreover, attentional selection appears to rely on different neural mechanisms depending on the structure of the scene: in scenes where the overlapping sounds are from the same category, attention primarily enhances object-level representations, whereas in cross-category scenes it engages category-level distinctions^25,26^. Thus, attention may flexibly target both object- and feature-level representations, depending on which features provide the greatest contrast between overlapping sounds^21,27^. However, this hypothesis has yet to be tested by directly quantifying the relative contributions of low-level and abstract sound features to attentional modulation across diverse soundscapes.

Another challenge in studying the neural basis of auditory processing is that different sound features are inherently correlated with each other^1,28^. Hence, to dissociate the role each sound feature plays in neural activity, the covariance between sound features must be considered. This problem has been partially solved in prior research by selecting or creating auditory stimuli with matched low-level profiles across abstract categories^12,29^. While such methods help isolate higher-level effects, they also reduce the ecological validity of the sounds, as they are by design highly similar in their low-level features. Another approach involves using statistical methods to remove covariances between models of different sound features^17,28,30^. This strategy has, however, not yet been adopted to delineate which sound features are modulated by attention in category-heterogenous scenes.

The aim of this study was to delineate the level of attentional modulation (low-level acoustic features/abstract features) in complex scenes containing natural sounds. For this, we analysed data from an fMRI paradigm^25^ where participants were presented with sounds from three natural sound categories: speech, instrument sounds, and animal sounds in three separate experiments (Figure 1). In the One Object Alone (1OA) experiment participants were asked to attend to one sound, which was played in isolation. In the three objects across categories (3OA) experiment, participants selectively attended to one sound from a soundscape which contained a sound from each of three different sound categories (e.g., Man, Trumpet, Bird). And finally, in the three objects within category (3OW) experiment, participants selectively attended to one sound from a soundscape consisting of three sounds from the same category (e.g., Bird, Monkey, Whale). We used representational similarity analysis (RSA)^31,32^ to model low-level and abstract sound features (Figure 2) and removed covariance between the feature models^28^. We then computed representational similarities between sound-feature models and neural activation patterns. Attentional modulation of a given sound feature was quantified as the difference in its representational strength between attended and distractor conditions (Figure 3). We finally used cross-experiment decoding to evaluate separately for each category which brain regions contain information that distinguish different objects of the category and whether attention boosts such representations (Figure 4).

**Figure 1.**
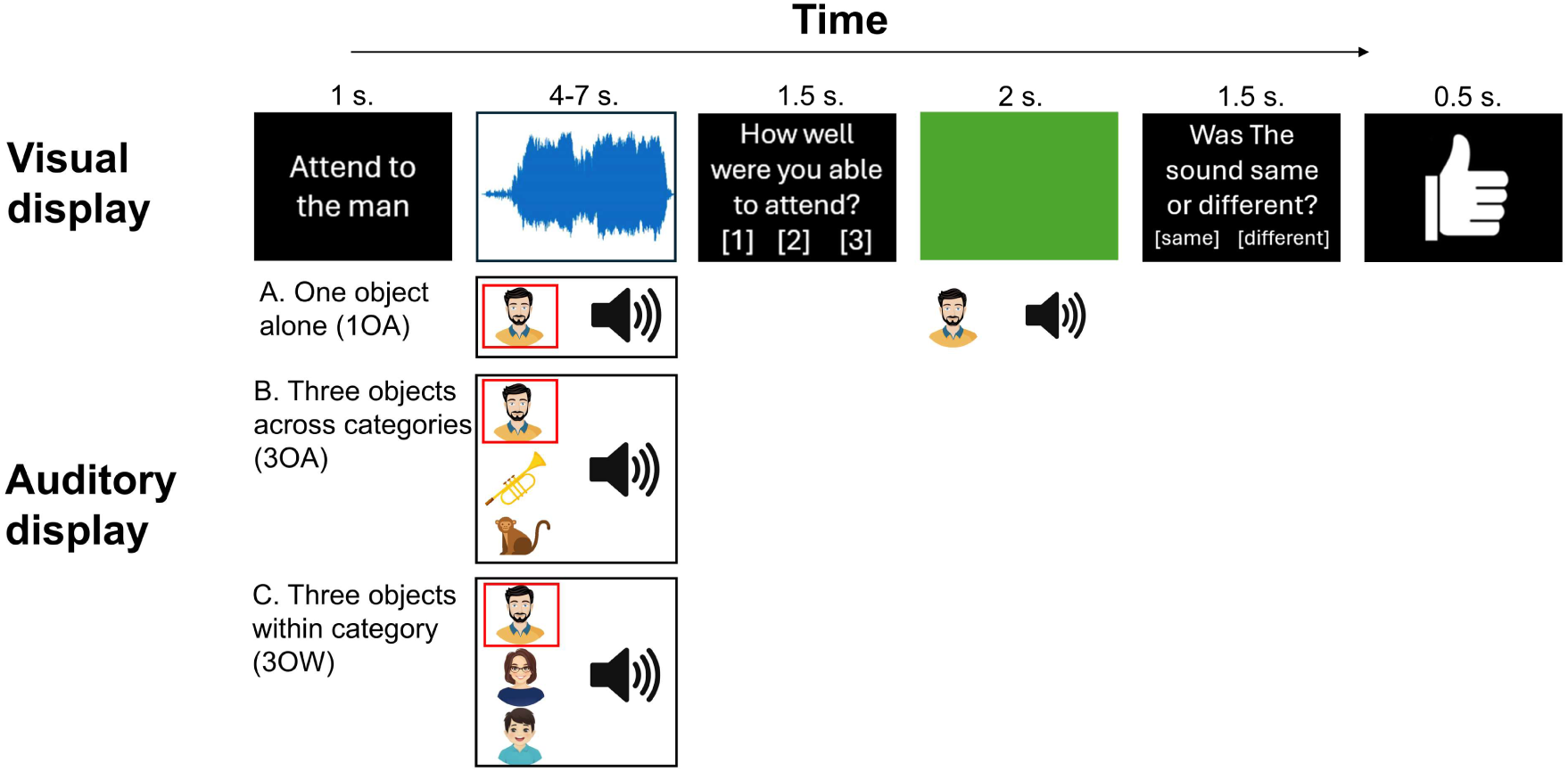
A schematic representation of the trial structure and experiments. Trials started with an instruction which oriented the participant to attend to a specific subcategory (e.g., Male1). After the instruction the participant was presented with stimuli. A) In the One object alone (1OA) experiment, the participant was presented with only one auditory stimulus. B) In the three objects across categories (3OA) experiment the participant was presented with one auditory stimulus from each of the three categories. C) In the three objects within category (3OW) experiment the participant was presented with three auditory stimuli from the same category. The visual stimulus was also presented regardless of the experiment. After the stimulus presentation, the participant rated how well they were able to selectively attend to the target stimulus. In 25% of the trials participants completed a match-to-sample task. In the match-to-sample task the participant was presented with a short sample of either the target stimulus or another stimulus from the same subcategory. Afterwards the participant determined whether the sample was from the target stimulus or from a different stimulus. Finally, the participant was given visual feedback on whether the match-to-sample task was answered correctly or incorrectly.

**Figure 2.**
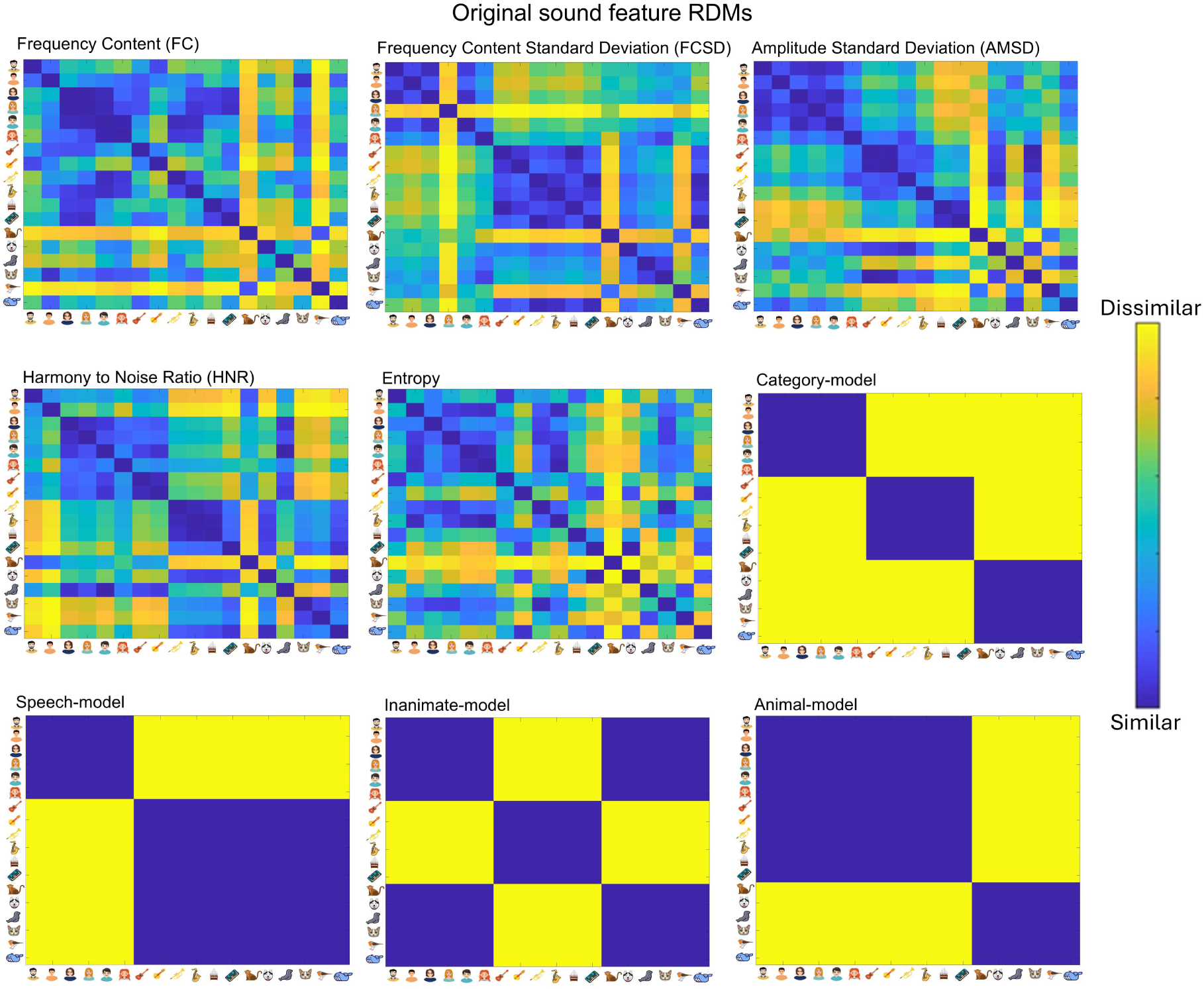
Original sound feature representational dissimilarity matrices. Here displayed are all 9 *Original* RDMs. Yellow colours indicate high dissimilarity between two subcategories for the given feature, whereas blue colours indicate high similarity.

**Figure 3.**
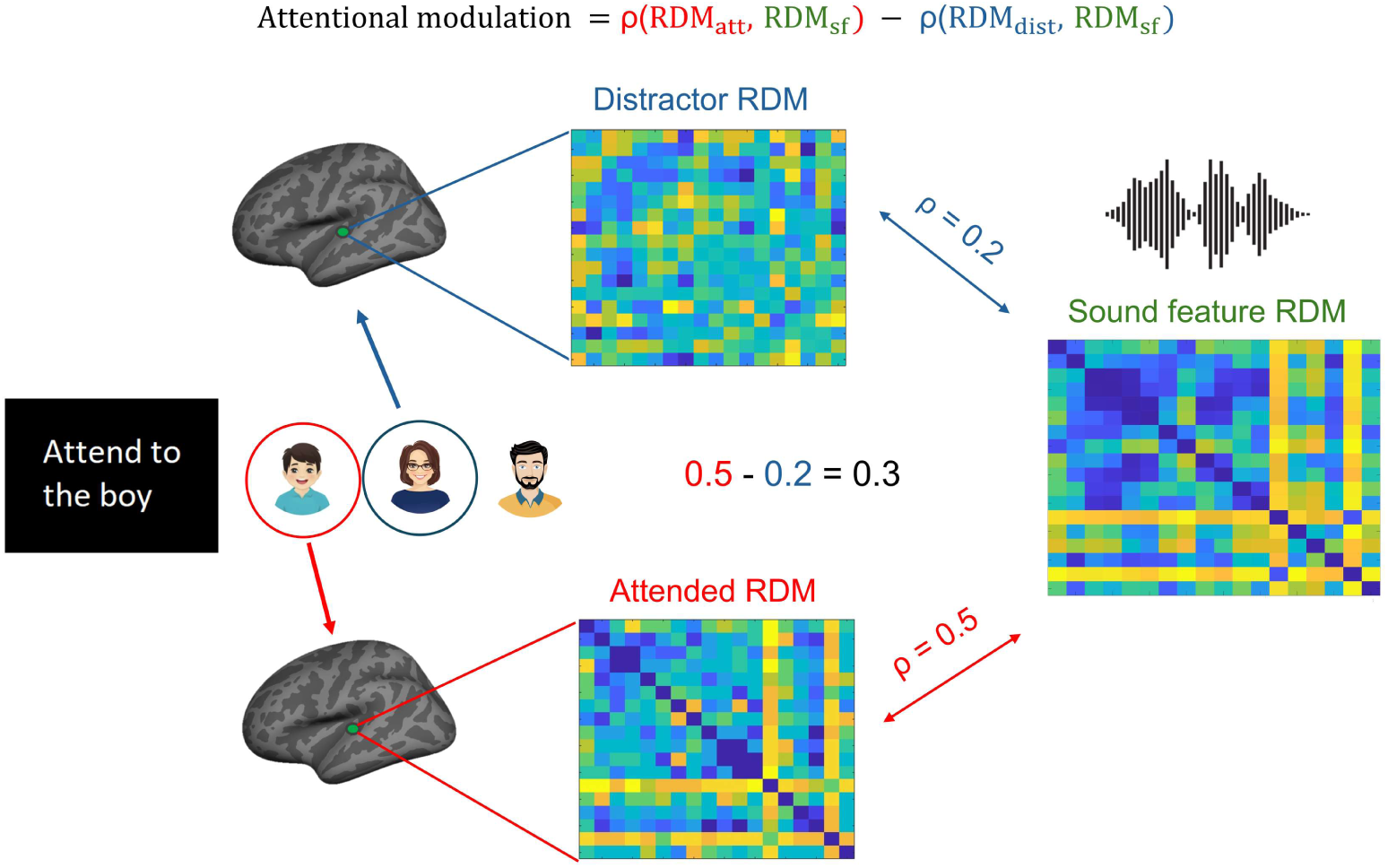
Schematic representation of RSA analysis pipeline for the selective attention tasks. Representational dissimilarity matrices (RDMs) were computed separately for attended and distractor sounds. Attended RDMs were generated by grouping trials with a common target sound into conditions and computing neural pattern similarities across these conditions. The same procedure was applied to distractor sounds to produce corresponding distractor RDMs. Sound feature RDMs capture the similarity structure of specific low-level or abstract sound features. Attentional modulation for each feature was quantified as the difference between its correlations to attended versus distractor RDMs. In other words, the attentional modulation index reflects how strongly the neural representational structure aligns with a given sound feature when attention is directed toward that sound compared to when it is ignored.

**Figure 4.**
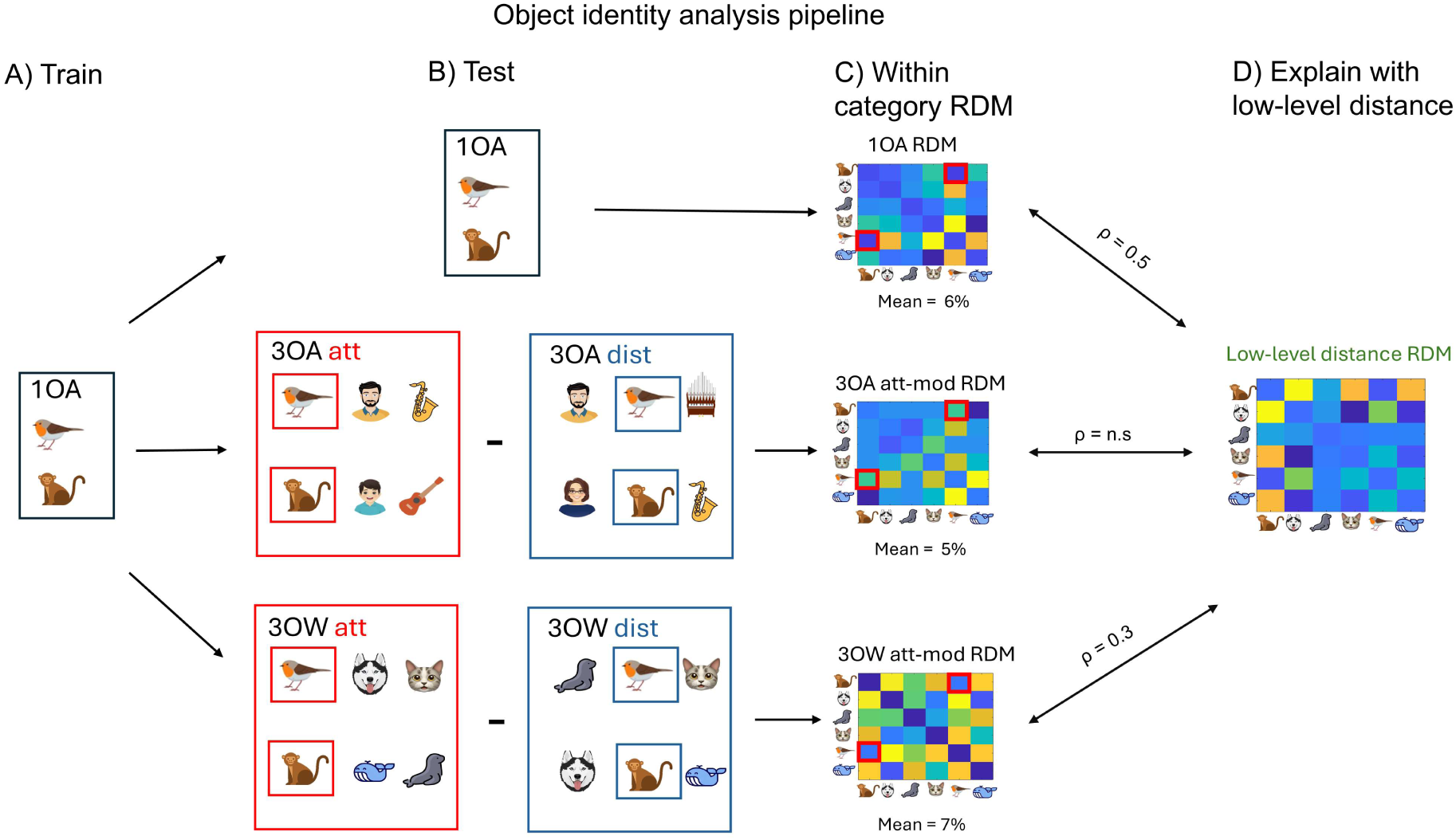
Cross-experiment classification analysis for object identity representations. A) A linear support vector machine (SVM) classifier was trained on trials from the one-object-alone (1OA) experiment, in which sounds from individual subcategories were listened to in isolation (e.g., Monkey vs. Bird trials). B) The trained classifier was tested on three conditions: held-out 1OA trials, selective-attention trials (3OA, 3OW) in which the same subcategories were attended, and trials in which the same subcategories acted as distractors. Classification performance was computed separately for attended and distractor conditions. C) Object-identity representation strength was defined as the mean classification accuracy across all unique within-category subcategory pairs. Attentional modulation was quantified as the difference between attended and distractor classification accuracies. D) To assess whether object-identity classification could be explained by low-level acoustic differences, a general low-level distance model was derived by computing the Euclidean distance of each object pair in the five-dimensional low-level distance space. Pairwise distances from this model were correlated with within-category decoding RDMs using Spearman correlation, testing whether classification accuracy was associated low-level differences.

In the listening task (1OA), robust representations of low-level sound features were observed in early auditory cortex, whereas abstract sound features were broadly represented across temporal and frontal regions. During selective attention, when sound objects could be distinguished based on category-level information (3OA), attention operated at an abstract category level rather than enhancing low-level feature processing. In contrast, when competing sounds belonged to the same category (3OW), representations of low-level sound features were selectively enhanced for attended sounds. Object-identity representations were modulated by attention in both types of sound scenes, in a category specific manner. These results highlight the dynamic and task-dependent nature of attention—i.e., attention modulates different levels of hierarchical processing based on which level provides the most distinct contrast between the overlapping sounds.

## Methods

### Participants

Twenty healthy native Finnish-speaking, right-handed individuals (8 females, age range 19–32 years, mean age 22.7, SD = 3.68) participated in the fMRI session. All participants had normal or corrected-to-normal vision and hearing, with no history of neurological or developmental disorders. To assess handedness, the Edinburgh Handedness inventory^33^ was used. Written consent was obtained from each participant prior to their involvement. For their time, participants received monetary compensation at a rate of 15 €/hour. The study received approval from the Ethical Review Board in the Humanities and Social and Behavioural Sciences at the University of Helsinki and was conducted in alignment with the Declaration of Helsinki.

### Stimuli

The experiment was run using Presentation software (version 24.0; Neurobehavioral Systems, Berkeley, CA, USA), which controlled both auditory and visual stimulation. Sounds were delivered binaurally via Sensimetrics S14 earphones (Sensimetrics, Malden, MA, USA), and the volume was individually adjusted for each participant at the start of the session to approximately 85 dB. Visual material was displayed on a screen and viewed through a mirror mounted on the head coil. Participants responded using an fMRI-compatible button box.

The auditory stimulus set consisted of sound clips from three object categories—speech, animal, and instrument sounds—each further divided into six subcategories (Table 1). Speech sounds were extracted from spoken dialogues recorded in a sound proof studio^34^. Instrument sounds were created from speech clips produced by the same talkers as the speech stimuli, with a different talker assigned to each instrument, such that the resulting instrument stimuli followed speech-like temporal patterns rather than forming musical sequences. Animal sounds were collected from free internet sources. Each subcategory included eight exemplars, yielding a total of 144 unique sound clips (3 categories × 6 subcategories × 8 exemplars). Clip durations varied between 4 and 7 seconds, with a mean length of 5.3 seconds.

**Table 1.** Sound categories and subcategories.

| Speech | Instrument sounds | Animal sounds |
| --- | --- | --- |
| Male1 | Baritone guitar | Monkey |
| Male2 | 12-string guitar | Husky |
| Female1 | Trumpet | Seal |
| Female2 | Saxophone | Cat |
| Boy | Grand organ | Chick |
| Girl | Bebop organ | Whale |

Visual stimuli were derived from the auditory stimuli used in the study. For each sound, we generated a scrolling waveform video in MATLAB, showing the waveform of the sound in blue on a white background (See Figure 1). The videos were presented independently of their corresponding auditory stimuli. The stimulus generation procedure is described in more detail in^25^.

### Trial structure

The trial structure (Figure 1) can be divided into four parts. 1.) In the beginning of each trial, the participant was visually instructed (1 sec.) to selectively attend to a specific stimulus (target stimulus). The target stimulus was either a specific auditory subcategory (e.g., “Attend to the Boy”) or the visual stimulus (“Attend to the video”).

2) Following the instruction, the participant was presented with experimental stimuli (4 – 7 sec.). The presented stimuli always included a visual stimulus and auditory stimuli. The auditory stimuli varied based on the experiment (See Experiments).

3) After the stimulus presentation, the participant was asked to rate how well they had been able to selectively attend to the target stimulus. The rating was conducted by pressing a button with an appropriate option: 1 (not at all), 2 (approximately 50 % of the time) or 3 (the whole time). The time window for the self-rating was 1.5 seconds long.

4) Lastly, to ensure that the participant attended to the target stimulus, on 25 % of trials where the target stimulus was auditory and on 100 % of the trials where the target stimulus was visual, the participant was asked to perform a match-to-sample task. To alert the participant of the match-to-sample task the background of the visual display was changed from black to green. Subsequently a short (2 sec.) additional stimulus was played. The additional stimulus was either a sample of the target stimulus (50 % probability) or a sample of another stimulus from the target subcategory (50 % probability). After the additional stimulus was played, the participants had to decide whether the additional stimulus was a sample from the target stimulus or from a different stimulus. The response was given by pressing one of two buttons (1.5 sec. response window). Results from behavioural data are reported elsewhere^25^.

### Experiments

The participants completed three experiments in total. The experiments differed in the presentation of auditory stimuli and the selection of the target stimulus. In the One Object Alone (1OA) experiment, only one auditory stimulus was presented during stimulus presentation, and the auditory stimulus always served as the target. Visual videos were presented concurrently but were never task-relevant, and participants were instructed to ignore them. In each 1OA trial, a different auditory stimulus was presented, allowing for the presentation of all 144 auditory stimuli in isolation (See Experimental session). The order of the stimulus presentation was pseudorandomised individually for each participant.

In the Three Objects Across categories (3OA) and Three Objects Within Category (3OW) experiments, three overlapping auditory stimuli were presented on each trial. In 3OA, the three auditory stimuli were drawn from different sound categories (e.g., Male1, Trumpet, Monkey), whereas in 3OW all three sounds belonged to the same category (e.g., Male1, Female1, Boy). On 75 % of trials, the target was an auditory stimulus. On these auditory-target trials, participants were instructed to selectively attend to the target sound while ignoring the other two sounds (distractor sounds) and the concurrently presented visual stimulus. To facilitate attentional orienting, the target sound began randomly from 250–500 ms before the onset of the distractor sounds. On the remaining 25 % of trials, the target was the visual stimulus. On these trials the participants were instructed to selectively attend to the video and ignore all auditory stimuli. The stimuli were pseudorandomized into trials individually for each participant. The pseudorandomisation was done in such a way that the following parameters were always fulfilled: 1) Each subcategory acted as the target stimulus twice and as a distractor 4 times per run. 2) No two trials used the same three auditory stimuli—i.e., each trial was acoustically unique. 3) All 144 experimental stimuli were used one time per run: 108 stimuli were presented during the auditory trials and 36 during the visual trials. Further details on stimulus pseudorandomisation are provided elsewhere^25^.

### Experimental session

Each run of the 1OA experiment included 36 trials and lasted approximately 6 minutes. In the 3OA and 3OW experiments, each run consisted of 36 auditory-target trials and 12 visual-target trials (48 trials in total), with a duration of approximately 8 minutes per run. For each experiment, participants completed 4 runs, resulting in 12 runs across the three experiments. All participants completed the 1OA task first. Thereafter, the order of the 3OA and 3OW experiments was counterbalanced across participants: half completed 3OA second and 3OW third, and the other half completed the experiments in the opposite order. Between runs, participants were given short breaks and an opportunity to talk with the researchers. After completing all functional runs, anatomical images were acquired. The total scanning session lasted approximately 100 minutes.

### Pre-trial

Participants practiced all experiments outside of the scanner before the session. Each experiment was practiced for two runs in total. Initially, each experiment was practiced with slower response times. After that, the experiments were practiced with the same pace as the actual paradigm. To mitigate potential learning effects, the stimuli used in the pre-trial were distinct from the stimuli used in the actual paradigm.

### Data acquisition

Functional imaging data were obtained using a 3T MAGNETOM Skyra whole-body scanner (Siemens Healthcare, Erlangen, Germany). A 32-channel head coil, with two channels removed to provide a clear view of the stimuli, was used. A T2* echo-planar imaging sequence was used to acquire 45 continuous oblique axial slices (TR = 1050 ms, TE 30 ms, flip angle = 75°, field of view = 21 cm, slice thickness = 2.7 mm, in-plane resolution = 2.7 mm × 2.7 mm × 2.7 mm). Each session included 12 runs: the first 4 runs (1OA experiment) produced approximately 360 volumes per run, while the final 8 runs (3OA & 3OW experiments) yielded about 570 volumes per run. In total, roughly 6000 volumes were collected per participant. After functional imaging, a high-resolution T1-weighted anatomical scan was performed (TE = 3.3 ms, TR = 2530 ms, voxel matrix = 256 × 256, in-plane resolution = 1 mm × 1 mm × 1 mm) for co-registration purposes.

### Sound feature analysis

All 144 main experimental auditory stimuli were analysed for low-level features using MATLAB R2025a. Each stimulus was characterized along five low-level features.

*Frequency content* (FC) describes how strongly different frequencies contribute to a sound. FC was obtained by applying a Fast Fourier Transform (FFT) to the sound’s waveform, yielding a vector of energy amplitudes across frequency bins.

All other features were calculated within 40 ms windows using a 20 ms step to capture temporal dynamics. For these features, window-wise values were summarised (mean/standard deviation) into a single value per feature and stimulus. Before feature computation, stereo signals were converted to mono, z-scored for amplitude normalization, and each frame was multiplied by a Hann window to reduce spectral edge artifacts. Low-energy frames were excluded using an amplitude threshold to prevent silent segments from biasing the estimates. The cut-off for a silent frame was defined as a per-frame root-mean-square (RMS) amplitude ≤ 0.0225 (amplitude was in std units).

*Frequency content standard deviation* (FCSD) indexes the temporal variability of a sound’s spectral profile. The spectral centroid for each window was calculated as the power-weighted average frequency, and FCSD was defined as the standard deviation of these centroid values across time (Supplementary Figure 1). *Amplitude standard deviation* (AMSD) captures variability in the sound’s amplitude across time. AMSD was derived by taking the standard deviation of the RMS amplitude across windows (Supplementary Figure 2). *Harmony to noise ratio* (HNR) characterizes the degree of periodicity in a sound, indicating how harmonic or noisy it is. For each window, a normalized autocorrelation was computed, the largest positive-lag peak within adaptive frequency bounds was identified, and HNR was defined as

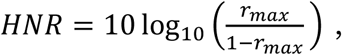

where *r*_max_ is the peak autocorrelation value. The resulting values were averaged across valid windows (Supplementary Figure 3). Spectral entropy quantified the average spectral complexity of each sound and was computed using the MIRtoolbox^35^ function mirentropy, then averaged across all valid windows (Supplementary Figure 4). The low-level sound features were selected based previous research^25,28,30^.

### Sound feature models

After computing the low-level sound features for each stimulus, we averaged the feature values within each subcategory (e.g., trumpet; Table 1). RDMs were then derived separately for each sound feature. For FC, which is a vector-valued feature (the magnitude spectrum across frequency bins), the RDM was constructed using 1 − *r*, where *r* is the Pearson correlation between the mean FC vectors of each pair of subcategories. For the remaining four scalar features (FCSD, AMSD, HNR, Entropy), each subcategory was described by a single scalar value, and the RDMs were constructed using the Euclidean distance between all pairs of subcategory means. In total, five low-level feature models were generated (Figure 2). To operationalize abstract sound features, four RDMs were constructed that divide the stimuli into categories. More specifically, we created a Category-model that equally differentiates each category, a Speech-model that differentiates speech from non-speech, an Inanimate-model that differentiates instrument sounds from animate sounds (animals and speech), and an Animal-model that differentiates animal sounds from non-animal sounds (Figure 2).

Next, to examine the covariance between the sound features, a Spearman correlation between each sound feature RDM was calculated (Supplementary Figure 5A). As can be seen from Supplementary Figure 5A, there exists substantial covariance between the sound feature RDMs. Without controlling for this covariance, it would be impossible to attribute any possible results to a specific sound feature. To address this issue, covariances between the models were statistically removed. First, RDMs were rank transformed to reduce sensitivity to scale differences and non-linearities in the dissimilarity structure. A general linear model was then used to explain shared variance between models, and the resulting residuals—capturing variance unique to each model—were retained for further analyses. These residualised RDMs are referred to as residualised models. Three sets of residualised models were created. For *Only Low-level models* all abstract RDMs were residualised out of each low-level RDM. This procedure eliminated correlations with abstract models while preserving correlations among the low-level features (Supplementary figure 5B). *Unique* models were obtained by residualising out all other models from each low-level feature model, thereby removing covariance with both abstract and other low-level models (Supplementary figure 5C). Importantly, some *Unique* models correlated only moderately with their original counterparts (the lowest being r = 0.73 between *Unique* FC and *Original* FC), indicating partial deviation from the original model structure. Finally, to create *Only abstract* models, all low-level models were residualised out of each abstract model, eliminating correlations between abstract and low-level models (Supplementary figure 5D). See supplementary figures 6 – 8 for all residualised models.

### Preprocessing and first-level fMRI data analysis

fMRI data was preprocessed using FEAT tool from FSL (FMRIB’s Sofware Library, http://www.fmrib.ox.ac.uk/fsl). Preprocessing included registration of fMRI volumes to the participant’s high-resolution structural image using FLIRT, motion correction using MCFLIRT^36^, slice timing correction (interleaved), non-brain removal using BET^37^ and high-pass temporal filtering using a cut-off of 100 seconds. The fMRI volumes were then transformed into the Freesurfer^38^ average surface space (fsaverage) using the Freesurfer function mri_vol2surfs, transforming the three-dimensional voxels into two dimensional vertices, without smoothing.

For the first level analyses, a GLM was fit to the time series of each vertex in each run. The GLM included a separate regressor for each trial. The GLM also included nuisance regressors for the 6 movement dimensions, to remove movement specific noise. This procedure resulted in trial specific spatial parametric maps.

### Multivariate pattern analyses

#### Representational similarity analysis

The fMRI RSA analyses were completed using both 8 mm searchlights^39^ with the help of Surfing toolbox^40^ as well as predefined regions of interest (ROIs; see Regions of interest). The first step of the fMRI RSA analysis was to make RDMs of neural activity. To do this, for each experiment separately, we grouped all trials in which participants listened (1OA) or attended to (3OA & 3OW) the same subcategory (e.g. trumpet) into conditions, resulting in 18 (one per subcategory) conditions per experiment. Each subcategory was listened to (1OA) or attended to (3OA, 3OW) eight times in total per experiment (two trials per run across four runs; see Experiments), yielding eight exemplars per condition.

Neural similarity between conditions was quantified using a linear support vector machine^41^ (SVM). The classifier was trained and tested with a leave-one-run-out cross-validation scheme: three runs served as training data (2 trials × 3 runs × 2 conditions = 12 exemplars for training), and one run was reserved for testing (2 trials × 1 run × 2 conditions = 4 exemplars for testing). This procedure was repeated four times so that each run was used once as the test set, and mean accuracy across folds represented the classification accuracy between each condition pair. Repeating this procedure for all possible condition pairs produced an RDM of decoding accuracies which index how dissimilar a given brain regions activation pattern was for different ***attended*** subcategories (e.g., how dissimilarly a region activated when *Male1* versus *Female1* was attended or listened to; Attended RDM).

For the selective attention experiments (3OA, 3OW), additional RDMs were created to capture the representational structure of distractor sound processing (See figure 3). The same decoding procedure was applied, but trials were grouped into conditions by using a shared distractor subcategory rather than attended subcategory. Because each auditory trial contained one target and two distractor sounds, the distractor conditions initially contained twice as many exemplars as the attended ones. To ensure comparability, the distractor trials were randomly restricted to an equal number of exemplars as the attended trials. The resulting RDMs reflect how dissimilarly cortical regions responded to different ***distractor*** subcategories (e.g., how dissimilarly a region activated when Male1 versus Female1 served as distractors; Distractor RDM).

Next, neural representations of sound features were derived. For the 1OA task this was done by computing Spearman correlations between sound feature models and the *Attended RDMs*. For selective attention tasks (3OA & 3OW), we also calculated Spearman correlations between the *distractor RDMs* and sound feature models. To isolate the effects of attentional modulation, these correlations were then subtracted from the corresponding correlations between *attended sound RDMs* and sound feature RDMs (Figure 3). This yields an attentional modulation index that quantifies the increase in representation of a sound feature when the sound is attended, relative to when it is ignored.

#### Object identity analysis

The analyses described above establish whether neural activity patterns represent low-level acoustic features and abstract category-level distinctions (e.g., speech vs. animal sounds). To further assess representations at a finer level of abstraction, we conducted a within-category differentiation analysis to determine whether neural patterns distinguish individual sound objects within a category—i.e., object identity (e.g., Male1 vs. Male2). This analysis was conducted separately for each category (speech, instrument sounds, animal sounds), to avoid category effects.

To index object identity representations, we implemented a cross-experiment classification scheme (Figure 4). A linear SVM classifier was first trained on trials from the One object alone (1OA) experiment, where sounds from a given subcategory were presented in isolation (e.g., Monkey vs. Bird trials; Figure 4A). The trained classifier was then tested under three conditions: 1) held-out 1OA trials, 2) trials from the selective-attention experiments in which the same subcategories were attended, and 3) trials in which those subcategories were present but acted as distractors (Figure 4B). For the held-out 1OA trials, classification followed the same leave-one-run-out cross-validation scheme described above (See Representational similarity analysis). In contrast, for cross-experiment decoding, the classifier was trained once using all 1OA trials for the given two subcategories and tested once on all corresponding 3OA/3OW trials in which that subcategory served either as the target sound or as a distractor. This procedure yielded 16 1OA exemplars for training and 16 3OA/3OW exemplars for testing per classification.

To quantify attentional modulation of object-level representations, we contrasted the classification accuracy between attended and distractor conditions in the selective-attention experiments (3OA and 3OW). That is, distractor classification accuracy was subtracted from attended classification accuracy to form an attentional modulation index. Positive values of this index indicate that neural patterns elicited by sounds in isolation (1OA) were closer to patterns observed during attended listening than to those observed when the same sounds were unattended. For each category, decoding accuracies were averaged across all unique within-category object pairs, and this mean value was interpreted as the strength of the object identity representation / attentional modulation (Figure 4C).

Finally, we examined whether the observed object-identity representations could be explained by systematic differences in low-level acoustic features. If object-pair classification accuracies were associated with low-level acoustic distance, this would indicate that classification performance is at least partly driven by low-level feature processing (or its attentional modulation), rather than abstract object-identity representations. To test this, we constructed a composite low-level acoustic feature RDM using Euclidean distance. Individual low-level feature RDMs (See sound feature models & Figure 2) were first scaled to unit variance without mean-centering, ensuring comparable feature scales while preserving the meaningful zero point of the RDM space (0 = no difference between objects for that feature). The scaled feature dimensions were then combined by computing the Euclidean distance from zero in the 5-dimensional low-level feature space for each object pair, yielding a general low-level acoustic distance RDM. Spearman correlations were computed between within-category classification RDMs and the corresponding low-level acoustic distance RDMs for each category (Figure 4D). This analysis was conducted within predefined ROIs (see Regions of Interest).

### Regions of interest

We used five ROIs for both hemispheres that were selected based on theoretical relevance (Table 3). These ROIs included key regions of the auditory ventral stream^3^. The ROIs were defined by combining subregions from the human connectome project atlas^42^ (see Table 2 & Figure 7).

**Table 2.** Selected ROIs and the corresponding ROIs from the human connectome atlas.

| Region of interest | Human connectome atlas regions of interest |
| --- | --- |
| Primary auditory cortex (A1) | A1 |
| Belt areas | MBelt + PBelt + LBelt |
| Superior temporal gyrus (STG) | A4 + A5 + TA2 + STGa |
| Superior temporal sulcus (STS) | STSda + STSdp + STSva + STSvp |
| Inferior frontal gyrus (IFG) | 45 |

### Inference of statistical significance

For the searchlight analysis, RSA-derived correlation (1OA) and attentional modulation (3OA/3OW) maps were first Fisher Z-transformed and then smoothed on the fsaverage cortical surface (FWHM = 5 mm). Group-level inference was subsequently performed using a cluster-forming threshold of Z = 3.1. Statistical significance in the searchlight analysis was assessed using cluster-based permutation testing (p < 0.05, corrected), as implemented in FreeSurfer’s *mri_glmfit-sim* tool (1,000 sign-flipping iterations). Only spatially contiguous clusters exceeding the cluster-size threshold derived from the null distribution were retained.

For the RSA ROI analysis, we tested whether the observed correlations/attentional modulations significantly differed from 0 using a two-sided t-test. P-values were corrected for multiple comparisons across all experiments sound features and ROIs using the Benjamini–Hochberg false discovery rate (FDR) procedure with α = 0.05.

The same approach was taken with the object identity analysis, where significant deviation from 0 for both the classification means and the low-level RDM correlations were tested using a two-sided t-test. P-values were corrected for multiple comparisons across all experiments and ROIs using the Benjamini–Hochberg false discovery rate (FDR) procedure with α = 0.05.

Finally, for the RSA ROI analysis, noise ceilings were computed separately for each ROI and experiment. For the 1OA experiment, noise ceilings were estimated using the standard RSA noise-ceiling procedure with Spearman correlations^43^. For the selective-attention experiments, we estimated noise ceilings for the attentional modulation index. For each ROI and experiment, we computed a group-level modulation template as the difference between the mean attended and mean distractor RDMs (mean over participants). For each subject, we then correlated both the attended and distractor RDMs with this template and defined the subject’s modulation as the difference between these correlations. Lower and upper bounds were obtained using leave-one-subject-out versus inclusive group templates, respectively.

## Results

### Sound feature representations during isolated listening

#### Whole-brain correlates of low-level sound features

In the 1OA experiment, participants listened to one sound at a time—i.e., the task required active listening rather than selective attention. Figure 5 illustrates correlations between low-level sound-feature RDMs and searchlight-based neural activation patterns during this task. As shown in Figure 5A, the *Original* low-level sound-feature models exhibited widespread correlations across temporal and frontal cortices, including higher-order regions such as the IFG.

**Figure 5.**
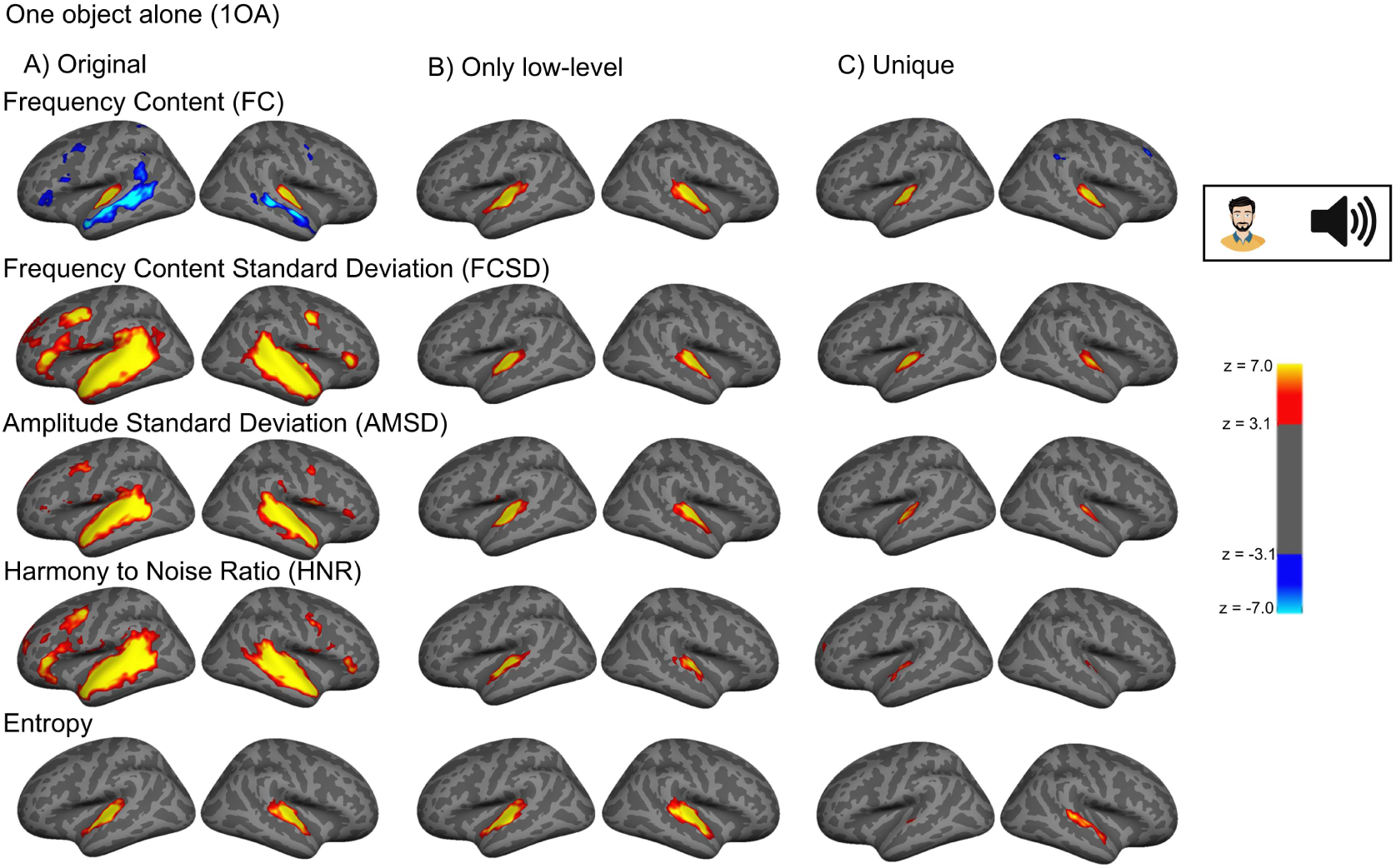
Correlates of low-level sound features during the 1OA experiment. This figure illustrates significant correlations between low-level sound feature RDMs (Figure 2) and searchlight-based activation pattern RDMs from the 1OA experiment. Brighter colours indicate stronger correlations, with warm colours indicating the presence of positive correlations and cold colours denoting negative correlations. Each row corresponds to a specific low-level sound feature, while each column represents a different version of the residualised models. A) column displays the *Original* models, B) column displays the *Only low-level* models, and C) column displays the *Unique* models. A schematic trial from the 1OA experiment is shown in the top right corner. Statistical overlays are displayed on FreeSurfer fsaverage inflated surfaces, where light gray indicates gyri and dark gray denotes sulci. Only lateral surfaces are shown here. All statistical parametric maps use an initial threshold of Z ≥ 3.1, with permutation-based cluster-size correction at p < 0.05.

Critically, however, when analyses were restricted to *Only low-level* sound-feature models (models that were statistically independent of abstract sound-feature models; see Supplementary Figure 5), these correlations were confined to auditory cortical regions. Across all low-level features, significant correlations were observed in A1, Belt areas, and STG. These results indicate that the apparent correlations between *Original* low-level sound features and higher-order brain regions in the were caused by shared variance with abstract sound features, rather than genuine representations of low-level acoustic information.

Finally, when using *Unique* models that were statistically independent of one another, different low-level sound feature correlations were more independently localized in the auditory cortex. Specifically, FC, FCSD, and AMSD correlated primarily with A1 and Belt areas, whereas HNR and entropy correlated with the left and right STG, respectively (Figure 5C).

Together, these results emphasize the necessity of covariance control between sound-feature models. Whereas *Original* models failed to dissociate low-level and abstract representations, and *Only low-level* models localized all low-level features mainly to overlapping regions, *Unique* models revealed meaningful spatial differentiation across the processing of different low-level features. Accordingly, all subsequent low-level results are reported using *Unique* models.

#### Whole-brain correlates of abstract sound features

*Original* abstract sound-feature RDMs showed strong correlations with neural activation patterns across widespread temporal and frontal regions (Figure 6A). Category and Speech models correlated broadly across temporal and frontal cortex, while Inanimate- and Animal-models showed widespread correlations encompassing STG, STS, IFG, and premotor cortex (PMC).

**Figure 6.**
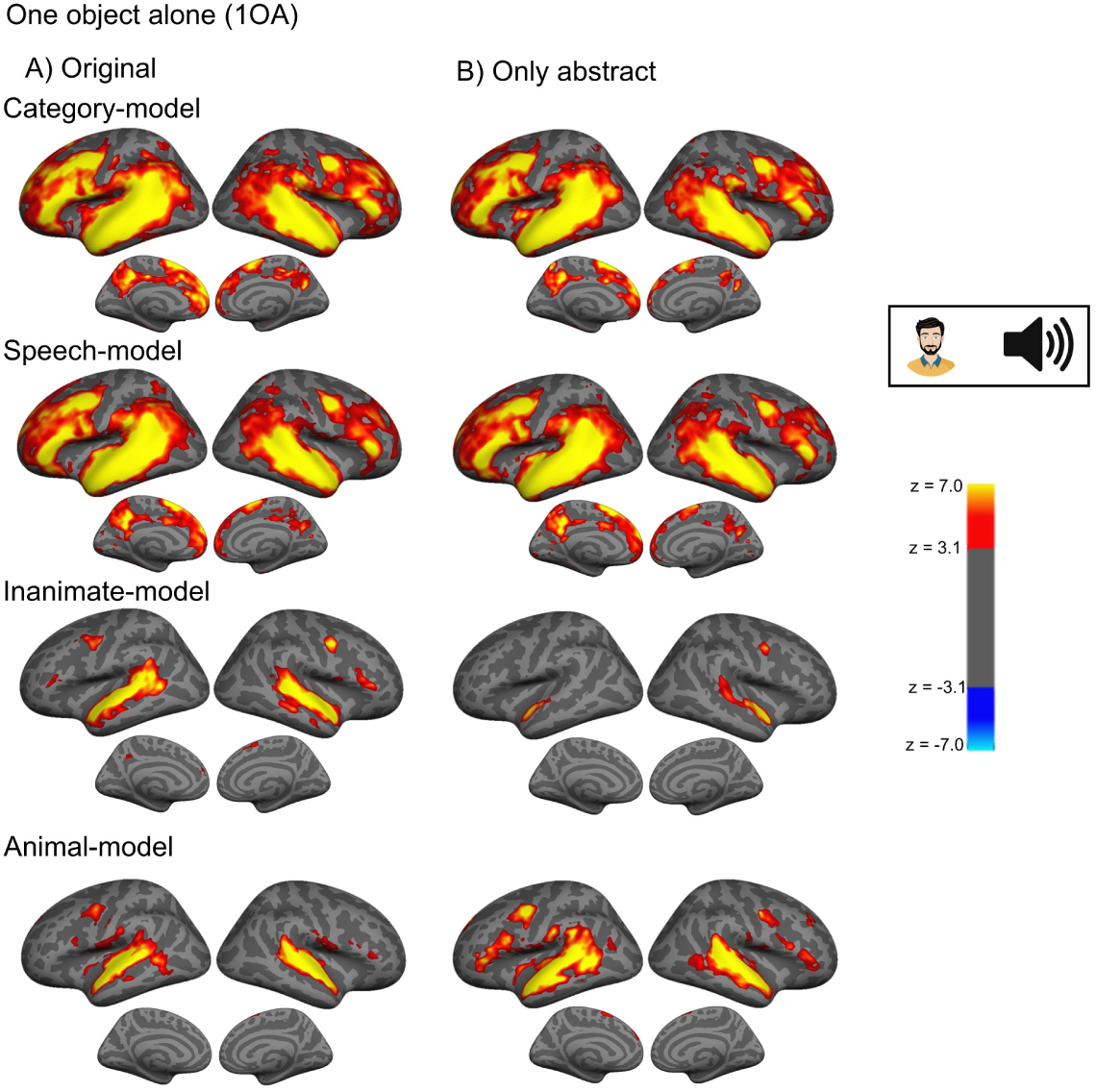
Correlates of abstract sound features during the 1OA experiment. This figure illustrates significant correlations between abstract sound feature RDMs (Figure 2) and searchlight-based activation pattern RDMs from the 1OA experiment. Rows correspond to different abstract sound feature models and columns represent different versions of the residualised models. Column A) shows results for the *Original* models, and Column B) shows results for the *Only abstract* models. Brighter colours indicate stronger correlations, with warm colours denoting positive correlations and cold colours denoting negative correlations. Lateral and medial surfaces are shown.

Using *Only abstract* models—defined as abstract feature models residualised with respect to low-level acoustic feature models—revealed several systematic changes in the observed correlations (Figure 6B). First, correlations for the Category-, Speech-, and Animal-models were modestly reduced in dorsal A1, consistent with this region’s primary sensitivity to low-level acoustic structure rather than abstract sound properties^5^. Second, correlations for the Inanimate model were markedly attenuated following residualisation and became largely confined to anterior STG. In contrast, correlations for the *Only abstract* Animal model were more spatially extensive than those observed for the Original model.

Notably, the differences between the Original and *Only abstract* Inanimate- and Animal-models were most pronounced in higher-order speech-selective regions, STS and IFG. Importantly, these regions did not exhibit significant correlations with low-level acoustic feature models (Figure 5C). Thus, although residualisation was intended to remove variance shared with low-level sound features, its effects were most evident in areas that were not themselves sensitive to those features.

This apparent discrepancy is likely explained by changes in covariance with the Speech-model, which showed strong correlations with these regions (STS & IFG). Specifically, residualisation reduced the correlation between the Inanimate and Speech models (from 0.06 to −0.08), while increasing the correlation between the Animal and Speech models (from 0.06 to 0.14; Supplementary Figure 5A, C). Therefore, the observed differences between Original and *Only abstract* models in these regions may reflect altered covariance with the Speech model, rather than the direct removal of low-level acoustic feature variance.

#### Progression of sound feature processing along the auditory ventral stream

To more closely examine the progression of sound feature representations, we conducted the same analysis as above within predefined ROIs (Figure 7, middle; table 2). For visualization purposes, we show mean RDMs across participants within each left hemisphere ROI from the 1OA experiment (Figure 7, top). As can be seen in Figure 7 top, activity patterns change along the ventral stream from initially dissociating monkey & bird sounds (acoustic outliers) to regions that distinguish speech sounds from other sounds (abstract feature). To test this qualitative observation, we correlated the ROI RDMs with the sound-feature RDMs (Figure 7A–B). Here, we report results for the *Unique* and *Only abstract* models, which maximize interpretability. Results for the *Original* and *Only low-level* models are provided in Supplementary Figure 9.

**Figure 7.**
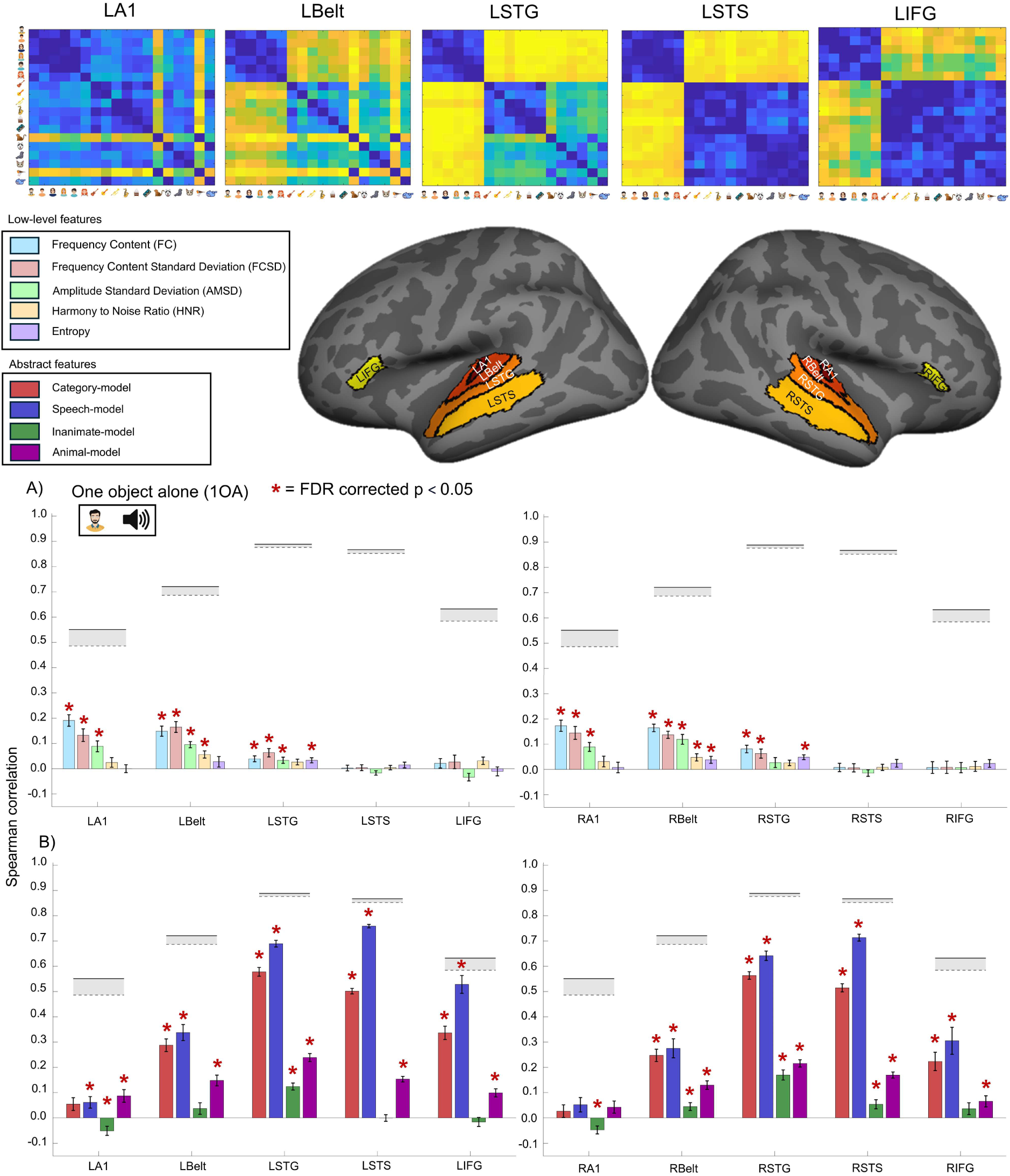
Region of interest analysis for the One object alone experiment. The top panel shows mean representational dissimilarity matrices (RDMs) from the 1OA experiment for each left-hemisphere ROI, illustrating activity patterns along the auditory ventral stream. ROI locations are shown on the cortical surfaces below the RDMs. The bottom panels present Spearman correlations between sound-feature RDMs and ROI RDMs. To maximize interpretability, results from the *Unique* and *Only abstract* models are shown. A) Panels display correlations for low-level sound features, and B) panels display correlations for abstract sound features. Left and right columns show left- and right-hemisphere ROIs, respectively. Red asterisks indicate correlations that significantly deviate from zero (FDR-corrected p < 0.05). Error bars represent ±1 SEM. Grey bars denote estimated noise ceilings, with dashed and solid lines indicating lower and upper bounds, respectively.

Low-level sound-feature models correlated bilaterally with activation patterns in A1, Belt, and STG (Figure 7A). In contrast, abstract sound-feature models showed strongest correlations in higher-order auditory regions, with bilateral STG, STS, and left IFG exhibiting particularly strong correlations (r > 0.5) with the Speech model (Figure 7B), indicating robust discrimination of speech sounds from other sound categories.

### Attentional modulation of sound feature processing

To examine attentional modulation of sound-feature processing, we applied the same analysis pipeline as in the listening task (1OA) separately to attended and distractor sounds during the selective-attention tasks (3OA & 3OW). Attentional modulation was quantified by subtracting distractor correlations from attended correlations, yielding an index of the extent to which sound-feature representations were enhanced when sound objects were attended, relative to when they were ignored (Figure 4).

#### Attentional modulation of sound feature processing in cross category soundscapes

In the 3OA experiment, participants attended to one sound in soundscapes comprising three sounds from different categories (e.g., Male1, Monkey, Trumpet). In the whole-brain analyses, apparent modulation of low-level sound features was observed only when using the *Original* models (Supplementary Figure 10A), which share variance with abstract feature models and are therefore unlikely to reflect genuine attentional modulation of low-level feature processing. In contrast, robust attentional modulation of abstract sound-feature representations was observed in higher-order temporal and frontal regions (Figure 8). These effects were most pronounced for the Category and Speech models, which showed significant modulation in the STG, STS, IFG, and premotor cortex (PMC)

**Figure 8.**
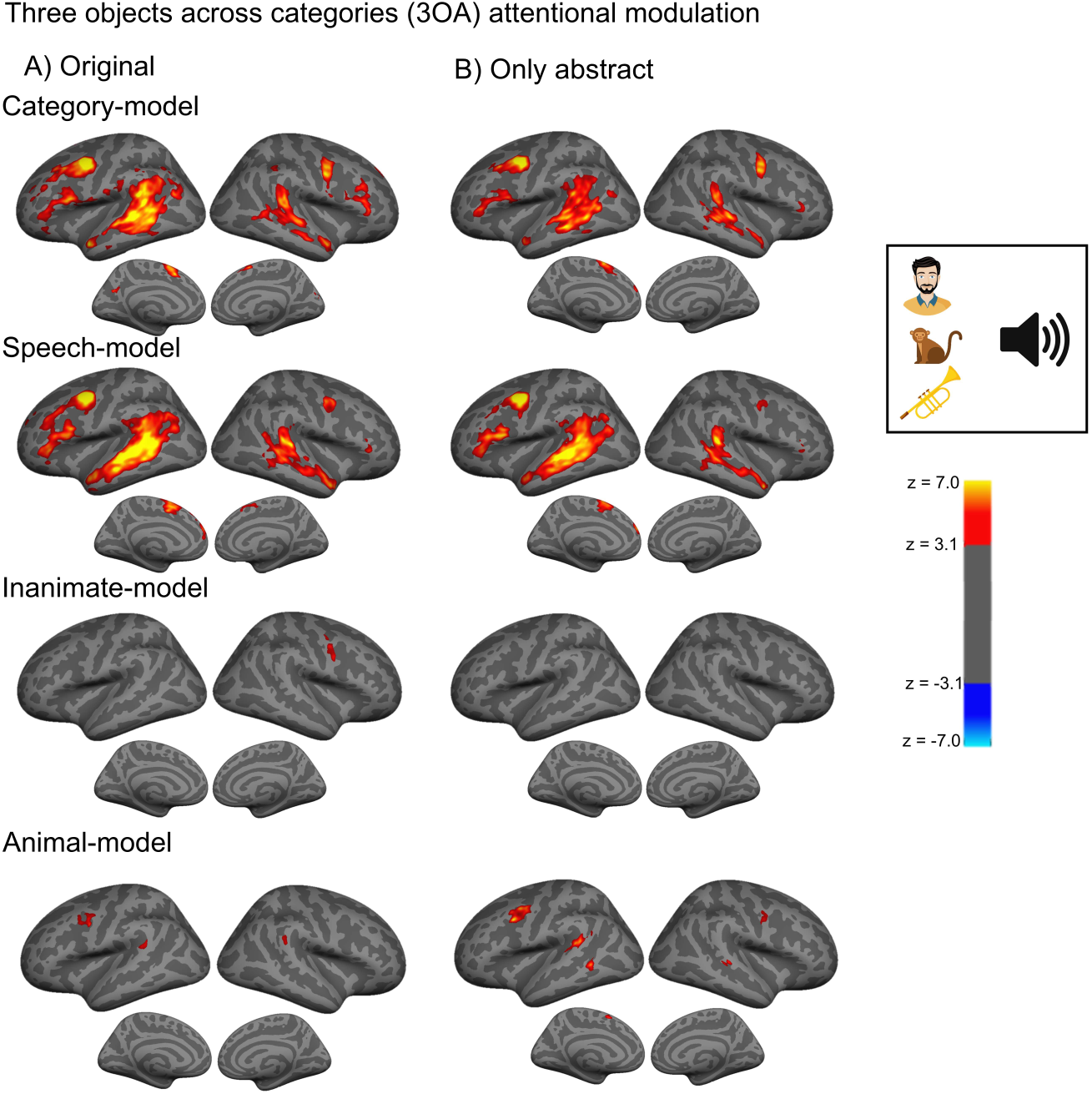
Whole-brain attentional modulation of abstract sound feature processing during the 3OA experiment. This figure shows searchlight-based attentional modulation effects for abstract sound feature models during the Three Objects Across Categories (3OA) task. Attentional modulation was quantified as the difference between correlations for attended and distractor sounds. Warm colours indicate stronger representations of attended relative to distractor sounds, whereas cold colours indicate the opposite. Rows correspond to different abstract sound feature models and columns represent different versions of the residualised models. Column A) shows results for the *Original* models, and Column B) shows results for the *Only abstract* models.

A similar pattern emerged in the ROI analyses. Attentional modulation of low-level sound features was again observed only for the *Original* models (Supplementary Figure 12A), but not for the Unique (Figure 9A) or *Only low-level* (Supplementary Figure 13A) models. In contrast, attentional modulation of abstract sound-feature processing was evident in STG, STS, and left IFG, with the strongest effects observed for the Speech model (Figure 9B). Together, these results indicate that during the 3OA task, attention did not modulate low-level acoustic feature processing, but instead selectively enhanced abstract, category-level representations.

**Figure 9.**
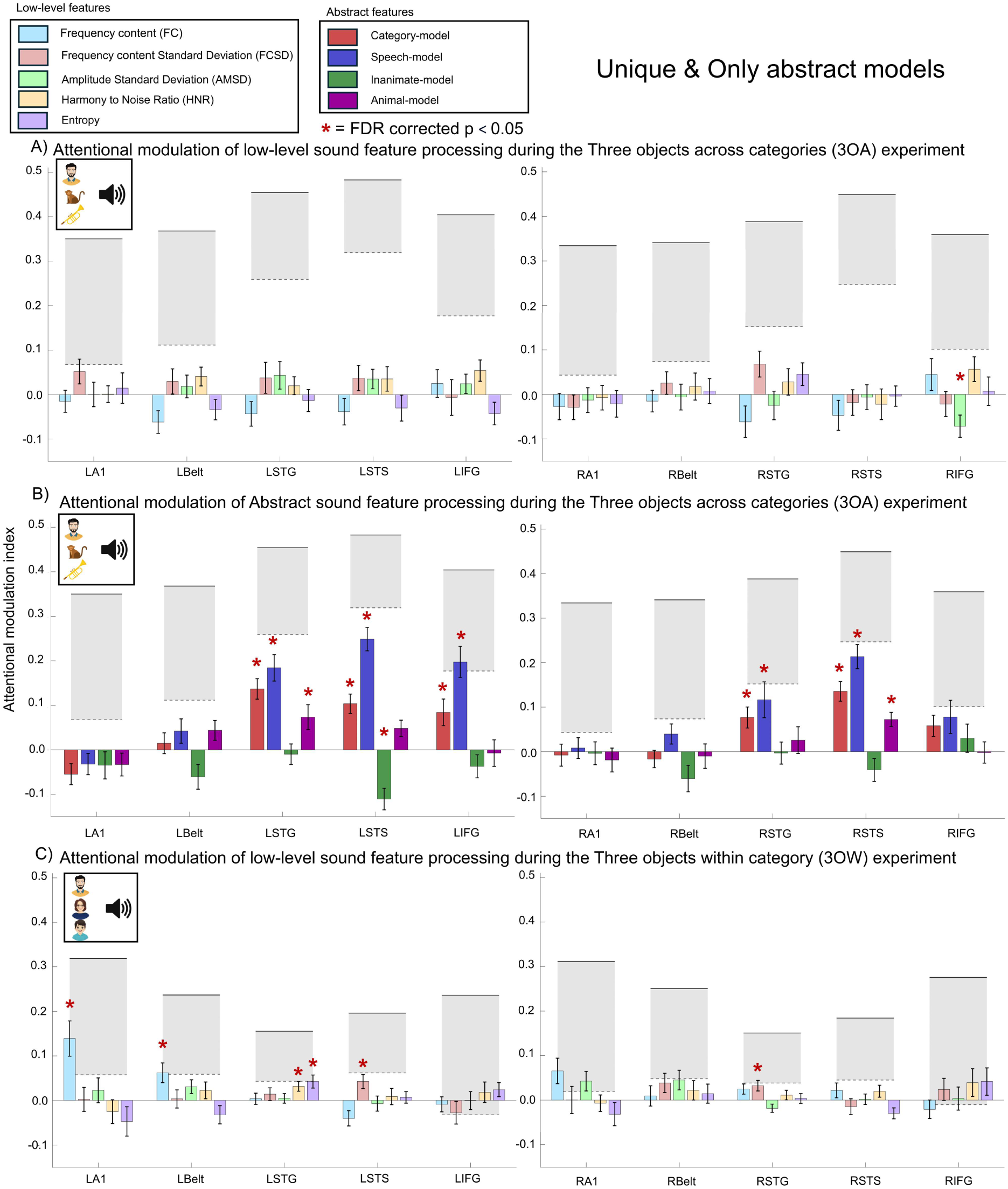
ROI analysis of attentional modulation of sound feature processing during selective attention. Attentional modulation indices (Figure 4) are shown for low-level and abstract sound features. To maximize interpretability, results are shown for *Unique* and *Only abstract* models. Panels A and B show results from the Three Objects Across Categories (3OA) experiment for low-level A) and abstract (B) sound features, respectively. Panel C shows results from the Three Objects Within Category (3OW) experiment for low-level sound features. Bars represent mean attentional modulation indices for each ROI, shown separately for left and right hemispheres. Red asterisks indicate attentional modulation values that significantly deviate from zero (FDR-corrected p < 0.05). Error bars represent ±1 SEM. Grey bars denote estimated noise ceilings, with dashed and solid lines indicating lower and upper bounds, respectively.

#### Attentional modulation of sound feature processing in within category soundscapes

In the 3OW experiment, participants attended to one sound in soundscapes comprising three sounds from the same category (e.g., Male1, Female1, Boy). Overall, the whole-brain searchlight analysis did not indicate strong attentional modulation of low-level sound features (except for an effect for FCSD in the right middle STG, see Supplementary Figure 11).

In contrast, the ROI analysis showed clear attentional modulation of low-level feature processing (Figure 9C). Attention modulated the processing of FC in the left A1 and Belt areas. HNR and Entropy showed attentional modulation in left STG, whereas attention modulated FCSD processing in right STG and left STS. However, unlike the other effects, the FCSD modulation in left STS was not present when using the *Original* or *Only low-level* models (Supplementary Figures 12C & 13C), suggesting that this effect likely reflects an artefact from covariance removal rather than modulation of a genuine low-level sound feature.

### Within-category differentiation of object identity

Above, we show that attention modulates abstract category-level, feature processing (e.g., is the attended sound Speech) to separate sounds from different categories in the 3OA task. Likewise, we show that attention modulates low-level feature processing (e.g., is the attended sound high in frequency content) to separate sounds from the same category in the 3OW experiment. Importantly, in the 3OW experiment, attended and distractor sounds always belonged to the same category, making category-level distinctions unavailable for separating the overlapping sounds. Under these conditions, attentional selection may instead operate at the level of another abstract feature: object identity (e.g., is the attended sound Male1).

To test this hypothesis, we implemented a cross-experiment decoding approach (Figure 4). A classifier was trained on 1OA trials, in which individual sounds were presented in isolation. The classification accuracy was first evaluated on held-out 1OA trials to establish baseline object-identity representations. The classifier was then applied to the selective-attention experiments, with classification accuracy assessed separately for attended and distractor sounds. This approach allowed us to test whether sound identity could be classified more accurately when the sounds were attended compared to when they were unattended. To determine whether any observed classification could be explained by low-level acoustic differences rather than abstract object identity, we additionally looked for associations between the classification accuracy and low-level acoustic distance (See Methods, Object identity analysis).

#### Decoding of object identity during isolated listening

Figure 10 shows the results of the object-identity decoding analysis. When sounds were presented in isolation (1OA), reliable object-identity decoding was observed across auditory regions. Classification accuracy was significantly above chance for speech and instrument sounds in A1, Belt and STG, whereas for animal sounds it was above chance in these regions as well as in right STS.

**Figure 10.**
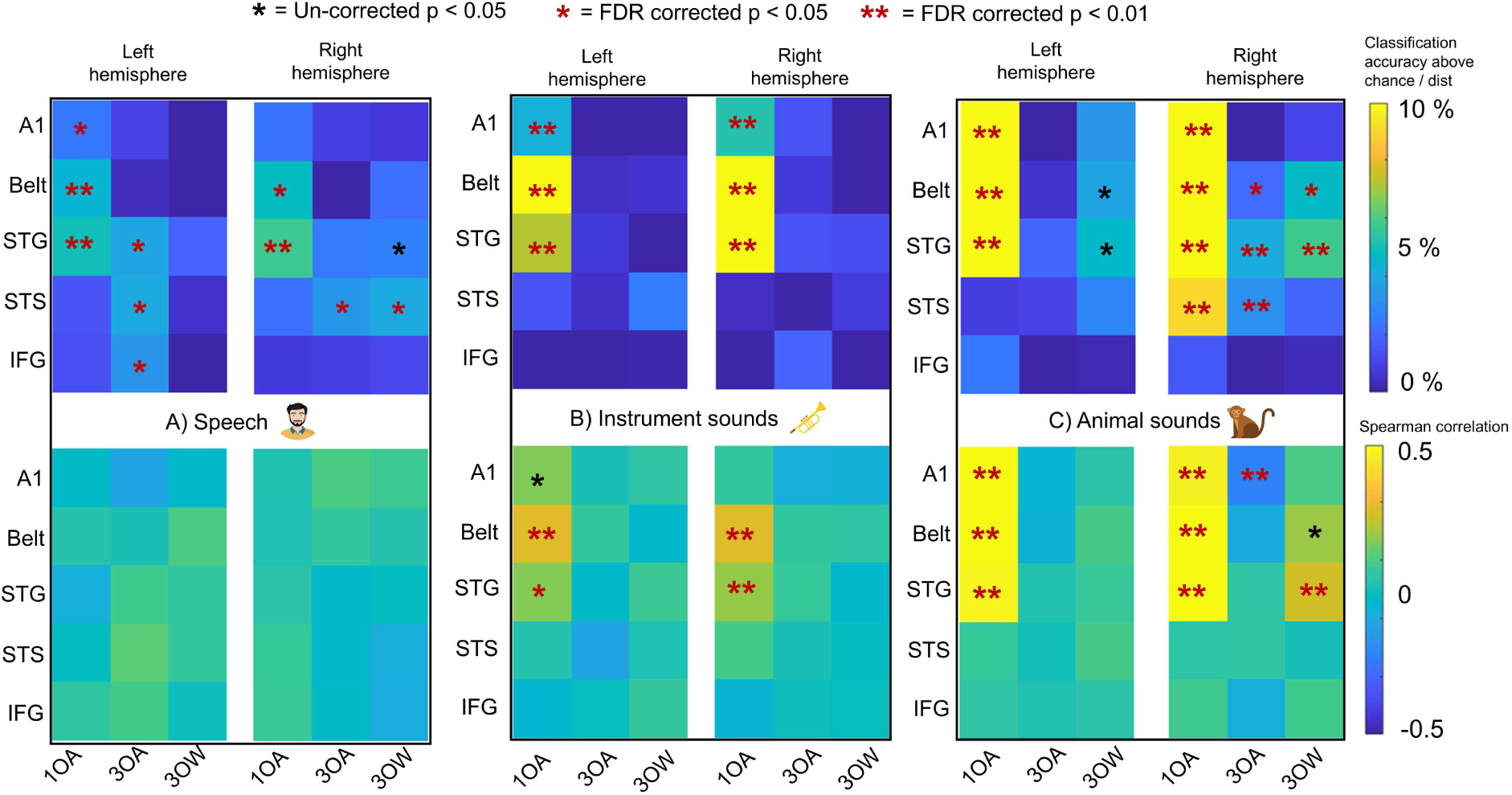
Decoding of object identity. Cross-experiment classification accuracies (top rows) are shown for speech A), instrument B), and animal C) sounds. The heatmaps display classification accuracies above chance for 1OA and above distractors (att > dist) for 3OA/3OW (in %). Bottom panel heatmaps show Spearman correlations between within-category decoding RDMs and low-level distance RDMs, assessing whether identity decoding can be explained by systematic acoustic differences. Asterisks indicate statistical significance (* uncorrected p < 0.05; red * FDR-corrected p < 0.05; red ** FDR-corrected p < 0.01).

To assess whether these identity representations reflected abstract identity rather than low-level acoustic separability, decoding performance was correlated with low-level acoustic distance measures. For instrument and animal sounds, decoding accuracy correlated significantly with low-level acoustic distance across most ROIs (with the exception of right A1 for instruments and right STS for animals), indicating that identity decoding in these categories can be at least partly explained by low-level sound feature differences. In contrast, no significant correlations with low-level acoustic distance were observed for speech sounds, suggesting that speech identity decoding was largely independent of low-level acoustic feature processing.

#### Attentional modulation of object identity processing

In the 3OA experiment, attentional modulation of object identity was observed for speech (Left STG, STS, IFG & Right STS) and animal sounds (Left Belt areas, STG & STS). These effects did not correlate with low-level acoustic distance, indicating that attentional modulation of identity representations in this experiment was not explained by attentional modulation of low-level sound features. This result is expected, as no attentional modulation of low-level feature processing was observed for this experiment (Figure 9A).

In the 3OW experiment, attentional modulation of object identity was observed for speech sounds in right STG and STS, and for animal sounds in left Belt and STG. Of these effects, however, only the speech effect in right STS survived FDR correction. Apparent attentional modulation of animal object identity was also observed in right Belt and STG, however, because these regions also exhibited significant correlations with low-level acoustic distance, these effects at least partly attributable to attentional modulation of low-level sound-feature processing, rather than reflecting pure object identity representations.

## Discussion

The present study investigated how selective attention modulates auditory representations at different levels of abstraction in complex soundscapes. By combining representational similarity analysis and cross-experiment decoding, we show that cortical attentional modulation is not fixed to a single representational stage. Instead, attention flexibly targets the representational level that is most informative for resolving competition, ranging from low-level acoustic features, abstract category-level features and object identity.

### Complexity of sound feature processing progresses along the auditory ventral stream

The 1OA experiment in this study closely resembled the experience of everyday listening to natural and isolated sounds and therefore allowed us to examine where different sound features are represented. The present study outlines five low-level sound features, which are processed at different levels of the auditory hierarchy independently of each other and the category of the natural sound being played. These findings are broadly consistent with previous work using similar acoustic features^30^, RSA-based methods^28^ and model free decompositions of auditory cortex activity^44,45^. Namely, a previous study identified six canonical response components^45^. The first two captured low- and high-frequency energy in A1, closely paralleling the FC-related representations observed here. Another component reflected sensitivity to harmonic pitch structure, with maximal weights near anterior belt areas and middle STG, consistent with the present HNR representations^45^. Interestingly, whereas previous work has suggested a coarse posterior–anterior dissociation between spectral and temporal sound feature processing^45,46^, our results indicate that spectral features (frequency content and its variability) and temporal features (amplitude variability) are represented in largely overlapping regions of primary auditory cortex (Figure 5C).

Together, these observations suggest a rough hierarchical organization of low-level sound feature processing. Features that require relatively little transformation of the acoustic signal—such as amplitude variability, frequency content, and frequency variability—were primarily represented in early auditory cortex. In contrast, less intuitive features such as harmonic strength and spectral entropy were predominantly represented in later regions, including belt areas and the STG, suggesting that these features may be used to bridge acoustic and semantic information^7^. Indeed, Prior work has implicated HNR as a second-order processing feature in natural sounds^47^, whereas, research in the visual domain speculates that image entropy may be used to reduce sensory redundancy, with strongest effects observed in non-primary visual cortex^48^, a pattern that appears homologous to the spectral entropy effects observed here.

Abstract sound features—most prominently the distinction between speech and non-speech—were represented broadly across temporal and frontal cortices (Figure 6), with particularly strong effects in STG, STS, and IFG, consistent with their established roles in speech processing^8–12,29^. In our previous work using univariate analyses on the same dataset, medial auditory cortical areas showed stronger responses to instrument and animal sounds, whereas speech elicited stronger responses in more lateral regions^25^. In the present study, we applied RSA to examine category-level representations and found that distinctions between sound categories were represented across both medial and lateral auditory cortex, as well as in frontal regions (Figure 6). Consistent with our earlier findings, however, an anterior region of the superior temporal gyrus (TA2) exhibited a particularly precise topography for instrument sound representations (Figure 6B), suggesting a degree of category-specific specialization within an otherwise distributed representational network.

Notably, both speech–non-speech and animal–non-animal distinctions were also decodable from the left A1 (Figure 7B), even after controlling for covariance with multiple low-level acoustic features. Given prior evidence that A1 exhibits task-dependent plasticity^49,50^, these effects may reflect category-dependent task demands that influence early sound encoding rather than genuine category abstraction. Support for this interpretation comes from the within-category object-identity analysis. Object-level decoding in A1 correlated with acoustic distance for animal and instrument sounds, but not for speech, indicating that different encoding strategies were engaged across categories (Figure 10). Whereas animal and instrument sounds appeared to be represented primarily via low-level acoustic features, speech sounds were more likely encoded at the level of object identity or linguistic information^51,52^.

Consistent with this view, activity patterns in primary auditory cortex are known to be sensitive to speech-specific articulatory structure^17,53^ and can be modulated by top-down influences arising from semantic context^21^. Additionally, prior work on non-speech human vocalizations has failed to observe robust within-category differentiation^28^, suggesting that the object-identity representations in early auditory cortex rely on speech-specific structure or linguistic relevance. The present findings are therefore consistent with the notion that speech may engage distinct encoding mechanisms already at early cortical stages. Another possibility is that apparent category-level representations in A1 reflect covariance with low-level features not explicitly modelled in the present study. Thus, although category-level decoding was observed in A1, these effects are unlikely to reflect category abstraction itself. Instead, they may arise from task-dependent encoding strategies or from residual acoustic structure not fully captured by the present feature models.

### Attentional modulations of sound feature representations are task-dependent

Does attention modulate early auditory processing to selectively represent low-level features of the attended sound? The present results suggest that it does, however, in a task-dependent manner. More specifically, in the 3OW experiment, attention selectively modulated FC in A1 and belt areas, and HNR, entropy and FCSD in the STG (Figure 7E). These findings replicate earlier demonstrations of frequency specific attentional modulation for simple sounds^13–16^, and extend them by showing that similar effects occur when selectively attending to natural sounds. Moreover, the results indicate that attentional modulation is not limited to the frequency content of the attended sound but can selectively enhance the processing of multiple independent low-level acoustic features. Importantly, attentional modulation extended beyond A1 to Belt areas and STG.

This result contrasts with several prior studies that failed to observe attentional modulation of low-level feature processing or early auditory cortex activity. One study found that BOLD responses in primary auditory areas were not explained significantly better by attended than unattended 10-min spoken narratives, whereas attentional modulation emerged in higher-order regions such as the superior temporal sulcus STS^19^. Using a similar paradigm with 10–15-min narratives presented alone or in overlapping pairs, another study reported attentional modulation already in primary auditory cortex, but the effect was selective for articulatory rather than spectral features^17^. Discrepancies in the results between these studies and the present work may provide insight into attentional modulation. First, attentional modulation of low-level features may be weaker for speech than for other sound categories. This interpretation is supported by our finding that apparent object-identity representations were at least partially explained by attentional modulation of low-level feature processing for animal sounds, whereas no such effect was observed for speech (Figure 10). This is probably because speech-specific features such as articulation^17^, semantic context^21^ or speaker identity^23^ may provide more efficient cues for selection. Second, attentional modulation of low-level feature processing may be amplified in soundscapes containing a larger number of overlapping sound objects—i.e., when sensory competition is increased^54^. Third, attentional modulation of low-level feature processing may be transient, such that during prolonged listening attention progressively shifts toward more abstract representations. This interpretation is consistent with spatiotemporal evidence showing that auditory attention initially modulates early sensory representations but rapidly reconfigures processing toward higher-order, object-level representations^55^ which may explain why sustained low-level modulation is difficult to detect during extended listening.

Electrophysiological work on speech^56–59^ and music^60^ has repeatedly shown that neural activity tracks the amplitude envelope of an attended stream more strongly than that of an unattended stream. Recent fMRI results have localized this effect to primary auditory cortex and the right medial STS^61^, implying that envelope tracking may depend partly on low-level sound-feature processing. Although stronger envelope tracking has been argued to reflect top-down semantic predictions rather than genuine low-level modulation^62^, the present findings show that attention can modulate low-level feature processing directly. This suggests that genuine low-level mechanisms may also contribute to enhanced tracking of attended speech. In the aforementioned fMRI study, envelope tracking was dissociated across cortical regions: primary auditory areas tracked the continuous amplitude envelope of the attended stream, whereas right STS tracked a more abstract, binary representation of speech presence (speaking vs not speaking)^61^. The authors proposed a division of labour in which primary auditory cortex encodes low-level stimulus features, while right STS represents higher-level, object-based aspects of the attended speaker^61^. The present results provide empirical support for this interpretation, as attentional modulation of low-level feature processing is observed in primary auditory cortex (Figure 9C), while the right STS expresses attentional modulation of speaker identity that is largely independent of low-level features (Figure 10A).

Importantly, modulation of low-level sound feature processing was not observed when the soundscape contained sounds from different categories (3OA). Instead, in those scenes, attention enhanced abstract category features represented in higher-level temporal and frontal regions with an especially strong effect in dissociating speech sounds from non-speech sounds^20^. This pattern suggests that auditory attention is adaptively guided by the representational level that best separates the competing sounds. When different sound categories are present, broad abstract distinctions help isolate the target, however when the sounds can’t be distinguished based on category, attention resorts to modulation of low-level sound feature processing.

### Within-category differentiation of sound objects depends on sound category

The object-identity analysis replicated previous findings showing that speaker identity can be decoded from neural activity patterns in the STG^51,52^. In addition, identity decoding for animal sounds was observed in right STS (Figure 10C), indicating that within-category object representations may not be restricted to speech but can also emerge for other sound categories.

Across both selective attention experiments (3OA & 3OW), we further observed attentional modulation of object-identity representations. In the 3OW condition, such modulation is expected, as selective processing over objects from the same category is required to resolve competition, and prior work has demonstrated attentional modulation of speaker identity in multi-speaker (category-homogenous) scenes^23^. However, the presence of attentional modulation in the 3OA condition is particularly informative, as it suggests that attention does not merely bias neural activity toward broad abstract category templates^25,26^ but also selectively enhances representations of individual objects within a category.

For speech identity representations a clear dissociation emerges: during isolated listening, speaker identity representations are expressed in lower-level auditory regions (A1, Belt, STG), whereas under selective attention, identity representations are selectively enhanced in higher-order regions (STG, STS, and IFG; Figure 10A). This pattern supports the notion that attention reweights representational processing toward higher-order regions that are better suited to resolving competition between sound objects^19^. A comparable shift was observed for the low-level HNR feature, which was primarily represented in Belt areas during isolated listening (Figure 7A), but showed attentional modulation in the STG under selective attention (Figure 9C).

Previous literature suggests a coarse lateralisation of speech processing so that the left hemisphere is concerned with semantic and linguistic information^2,63^, whereas the right hemisphere encodes speaker identity related information^9,51,52,64,65^. This lateralisation is partially supported here, as during the 3OW experiment, attentional modulation of speaker identity was observed exclusively in right-hemisphere STG and STS. In contrast, during the 3OA experiment, attentional modulation of speaker identity was left-lateralised to STG, STS, and IFG—regions that also exhibited attentional modulation of category-level representations (Figure 9B). Together, these findings suggest that the hemispheric lateralisation of speaker-identity processing may not be fixed, but depends on task demands and representational context, highlighting the need for future work to disentangle the specific mechanisms underlying lateralised identity representations.

Consistent with the pattern observed for object-identity representations during isolated listening, attentional modulation was not restricted to speech. Animal identity representations were also modulated by attention in both selective attention experiments (Figure 10C), suggesting that attentional modulation of object identity generalises to multiple sound categories. With animal sounds, attentional modulation of object-identity representation was observed in regions that exhibited acoustic dependence during isolated listening, specifically in Belt areas and STG (Figure 10C). This suggests that these regions may shift processing from predominantly acoustic encoding during isolated listening toward more abstract, object-level representations when attention is required to resolve competition among multiple sound objects. Within feature-integration frameworks, this pattern can be interpreted as attention promoting the integration of low-level features into coherent sound objects when selective processing is required^54,66,67^.

### Conclusions

This study investigated how low-level and abstract sound features are represented in the human brain during listening and selective attention using multivariate fMRI analyses. Three main conclusions emerge. 1) Low-level acoustic features of natural sounds were robustly represented in early auditory cortex, whereas abstract sound features were progressively encoded along the auditory ventral stream. This pattern is consistent with influential models proposing hierarchical processing of sound features with increasing representational abstraction along the ventral pathway^3^. 2) The level at which attention modulated auditory representations depended on how competing sound objects could be distinguished. When overlapping sounds could not be separated at the category level, attention enhanced low-level acoustic feature representations in early auditory regions. In contrast, when category cues provided sufficient separation, attentional modulation was primarily expressed at higher, more abstract representational levels. These findings demonstrate that auditory attention operates dynamically across hierarchical stages, selectively enhancing the representational level that provides the most effective contrast between competing sounds. 3) Beyond low-level and category-level effects, neural representations of object identity were observed. This effect was most robust for speaker identity, although some evidence suggests that it generalises to other sound categories. Importantly, attentional modulation of object-identity representations was observed even when category-level information was sufficient to distinguish competing sounds, indicating that attention can engage finer-grained object-identity representations beyond category-level processing.

## Author contributions

O.V.V.: Methodology, Software, Formal analysis, Investigation, Data curation, Writing—original draft, Visualization; I.A.M.: Methodology, Software, Formal analysis, Investigation, Data curation, Supervision, Writing—review and editing; P.A.W.: Methodology, Software, Investigation, Resources, Data curation, Funding acquisition, Supervision, Writing—review and editing.

## Acknowledgment

P.W. discloses support for the research of this work from the Research Council of Finland (grant numbers 1348353 and 368974). The funders had no role in the study design; data collection, analysis or interpretation; the decision to publish; or the preparation of the manuscript. We thank M. Salmikivi, W. Vikatmaa, S. Rossow and L. Lehtimäki for assistance with data acquisition; V. Laaksonen and J. Kauramäki for input on study design; and C. Pedersini for comments on the manuscript.

## Data Availability Statement

All preprocessed fMRI data, anonymized to protect participant privacy, will be deposited on GitHub: https://github.com/UHVaris/Attentional-modulation-of-sound-features

## Code availability

The custom code will be available in the GitHub repository: https://github.com/UHVaris/Attentional-modulation-of-sound-features.

