## Supplementary material for "Attention modulates neural representations of acoustic, categorical, and identity features in a task-dependent manner"

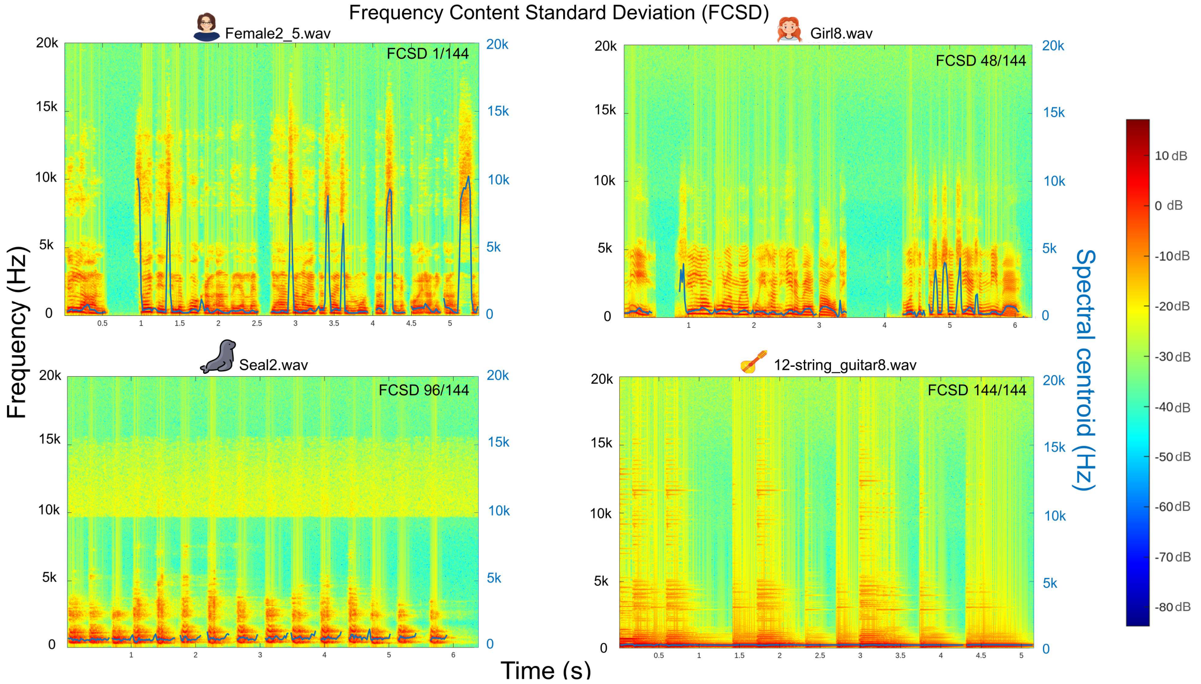


**Figure S1.** *Examples of Frequency content standard deviation (FCSD) calculation.* Spectrograms are shown for four stimuli representing the largest FCSD value, the 48th-largest, the 96th-largest, and the smallest FCSD value in the stimulus set. The estimated spectral centroid for each 40-ms frame is overlaid in blue. FCSD was defined as the standard deviation of the spectral-centroid trajectory over time. The stimulus with the highest FCSD value (Female2_5.wav) has large variation in the spectral centroid because it contains many “s” sounds which are very high in frequency.

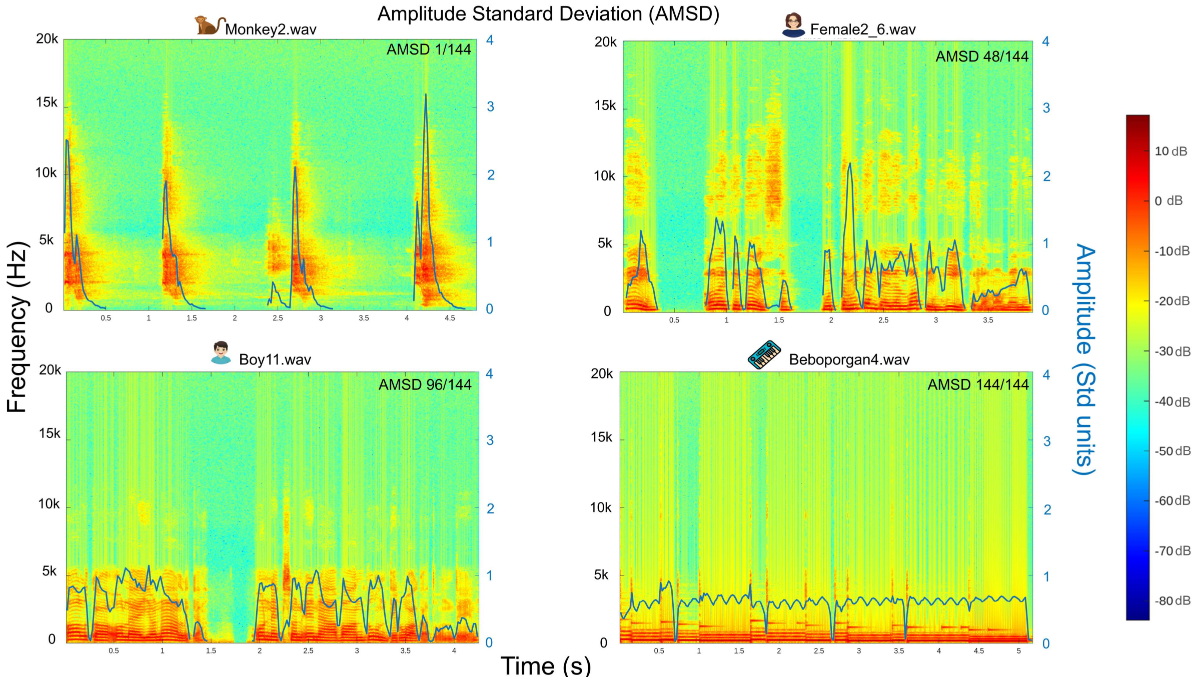


**Figure S2.** *Examples of Amplitude standard deviation (AMSD) calculation.* Spectrograms are shown for four stimuli representing the largest AMSD value, the 48th-largest, the 96th-largest, and the smallest AMSD value in the stimulus set. The estimated amplitude in standard deviation units for each

40-ms frame is overlaid in blue. AMSD was defined as the standard deviation of the amplitude trajectory over time.

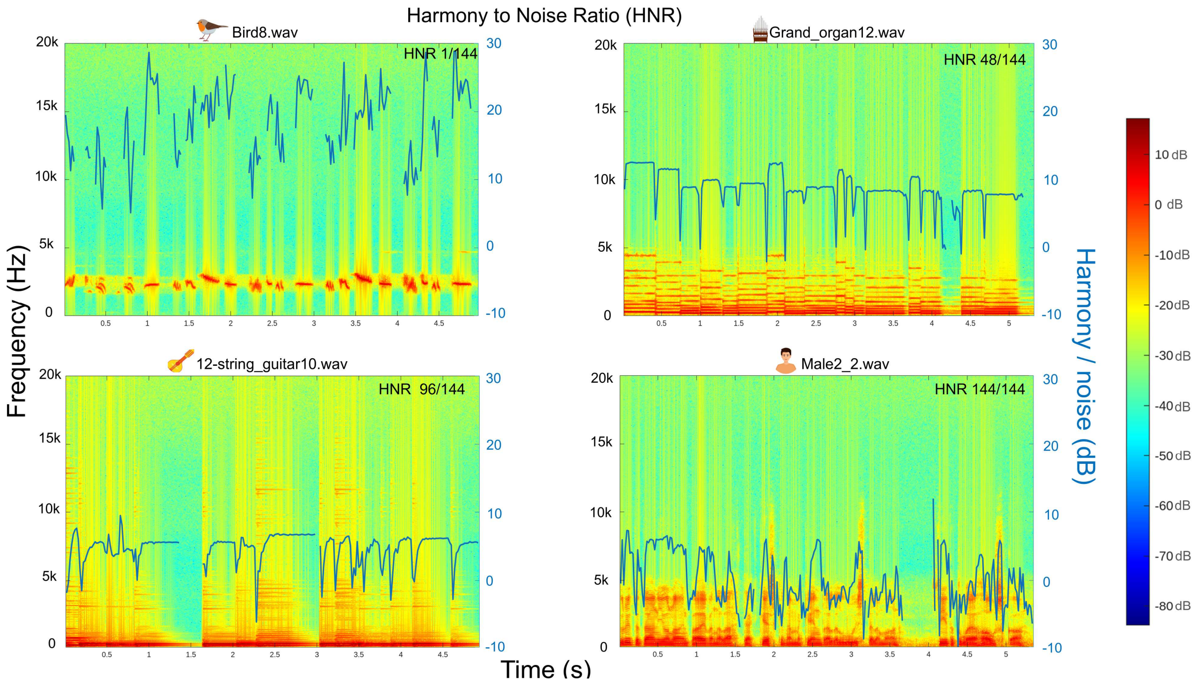


**Figure S3.** *Examples of Harmonics to noise ratio (HNR) calculation.* Spectrograms are shown for four stimuli representing the largest HNR value, the 48th-largest, the 96th-largest, and the smallest HNR value in the stimulus set. The estimated Harmonics to noise ration in dB units for each 40-ms frame is overlaid in blue. HNR was defined as the mean HNR value across time.

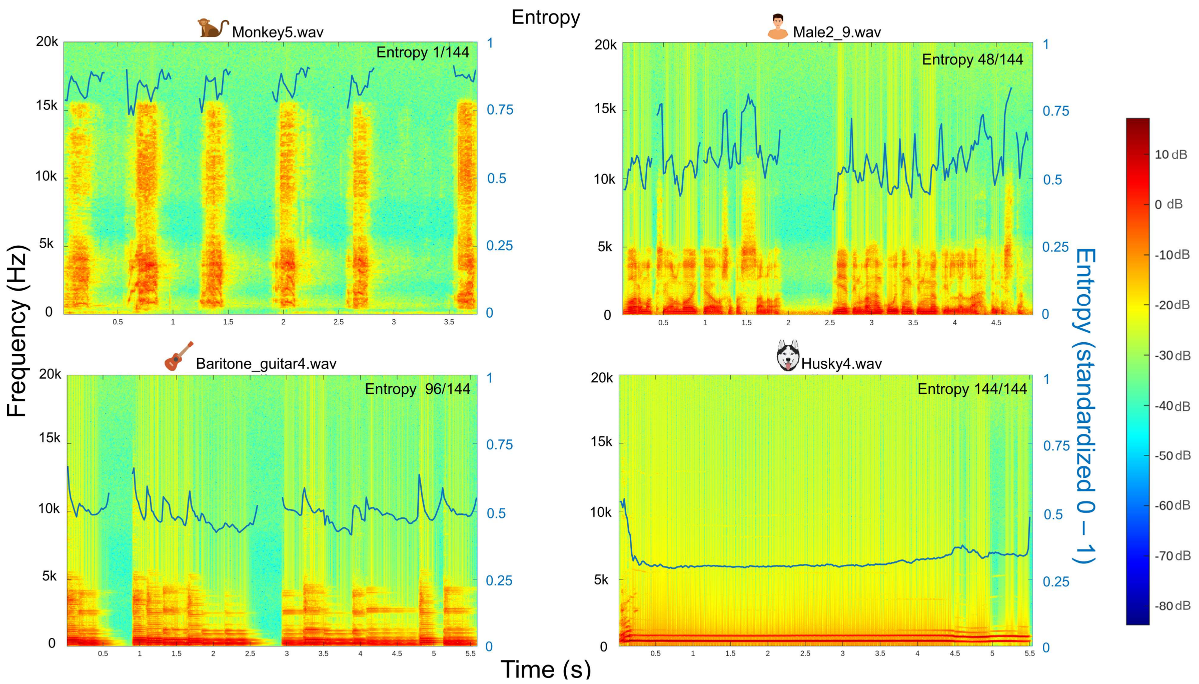


**Figure S4.** *Examples of Entropy calculation.* Spectrograms are shown for four stimuli representing the largest Entropy value, the 48th-largest, the 96th-largest, and the smallest Entrpy value in the stimulus set. The estimated entropy for each 40-ms frame is overlaid in blue. Entropy was defined as the mean entropy value across time.

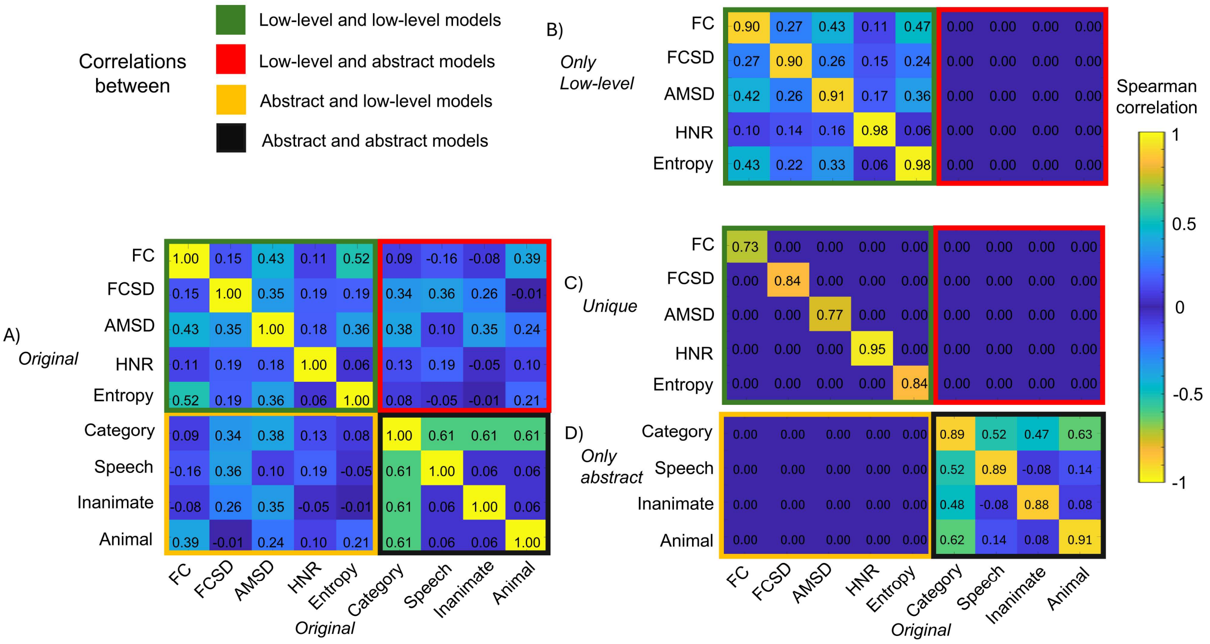


**Figure S5.** *Correlations between sound feature models*. Here displayed are Spearman correlation matrices between residualised models and original models. Bright and yellow colours indicate a strong correlation, whereas dark blue colours indicate a low correlation. A) Displays correlations between all *Original* sound feature models (Figure 2). Because the *Original* models are correlated against each other, the matrix is symmetric across the diagonal. The diagonal cells also have a correlation of 1, as the correlations are being calculated between the same models. B) Displays correlations between *Only Low-level* models and *Original* models. The *Only Low-level* models have no correlations with the *Original* abstract models (red box), as their covariance has been residualised out. C) Displays the correlations between the *Unique models* and *Original models*. As covariance with all other models has been regressed out, the *Unique models* show no correlations with any other *Original models*. D) Displays the correlations between *Only abstract models* and *Original models*. The correlations between *Only abstract models* and low-level models are 0, as they have been regressed out (orange box).

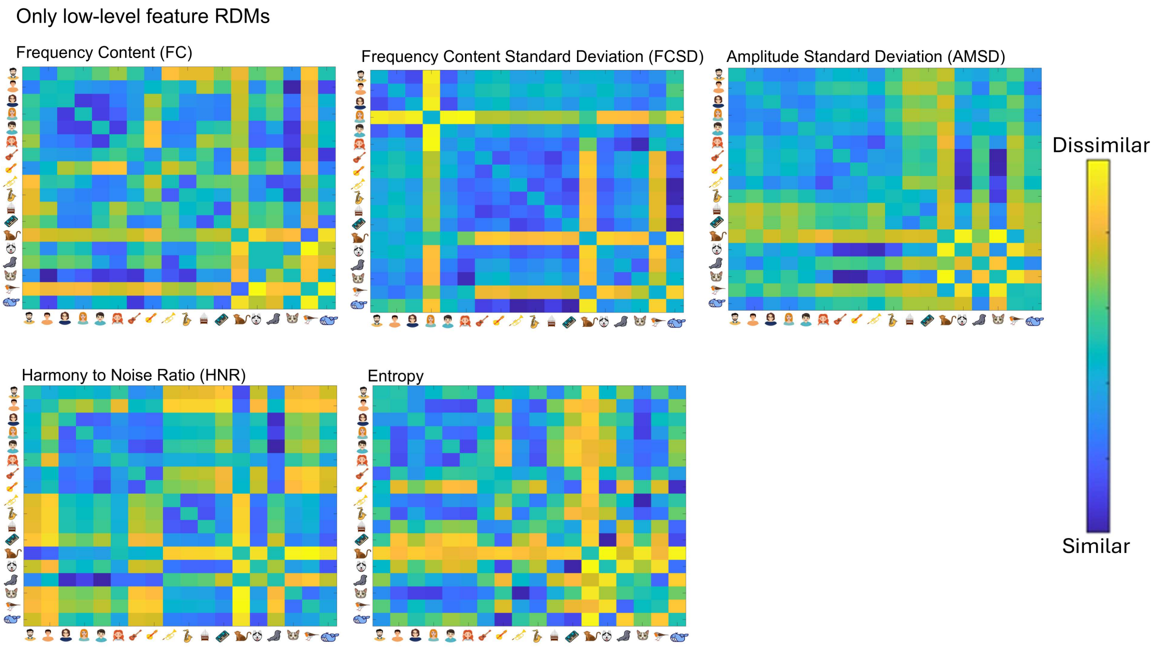


**Figure S6**. *Only low-level sound feature representational dissimilarity Matrices*. Here displayed are all *Only low-level* sound feature RDMs. These RDMs are independent of category level abstract sound features. Yellow colours indicate high dissimilarity between two subcategories for the given feature, whereas blue colours indicate high similarity.

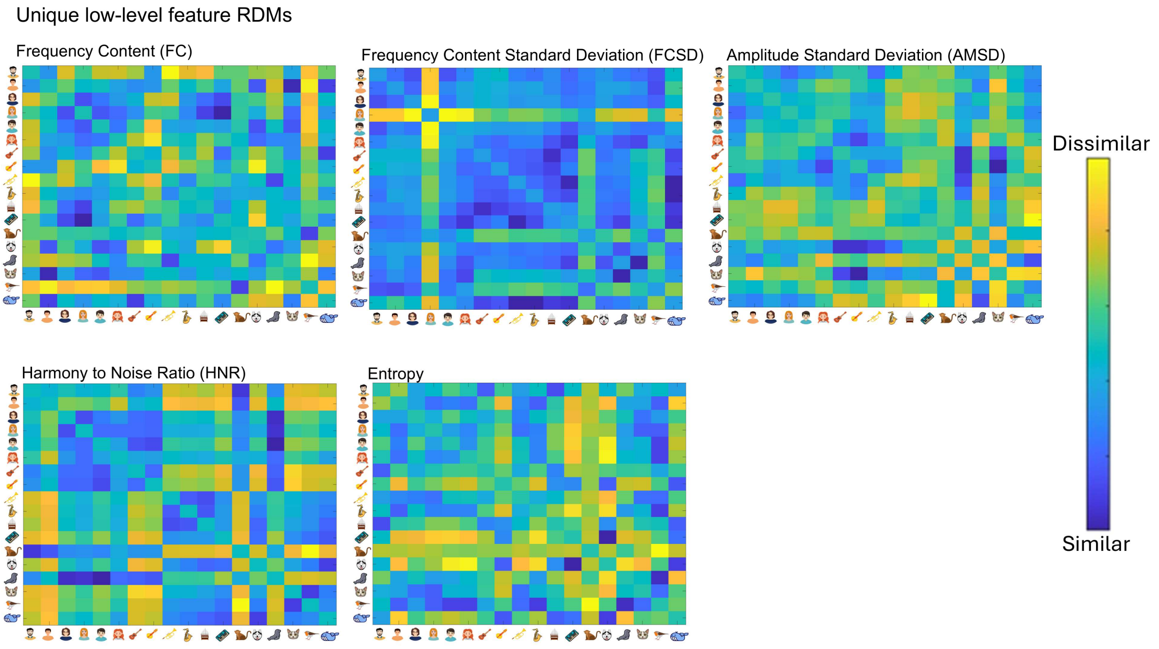


**Figure S7**. *Unique sound feature representational dissimilarity Matrices*. Here displayed are all 5 *Unique* sound feature RDMs. These RDMs are independent of all other sound feature models used in this study. Yellow colours indicate high dissimilarity between two subcategories for the given feature, whereas blue colours indicate high similarity.

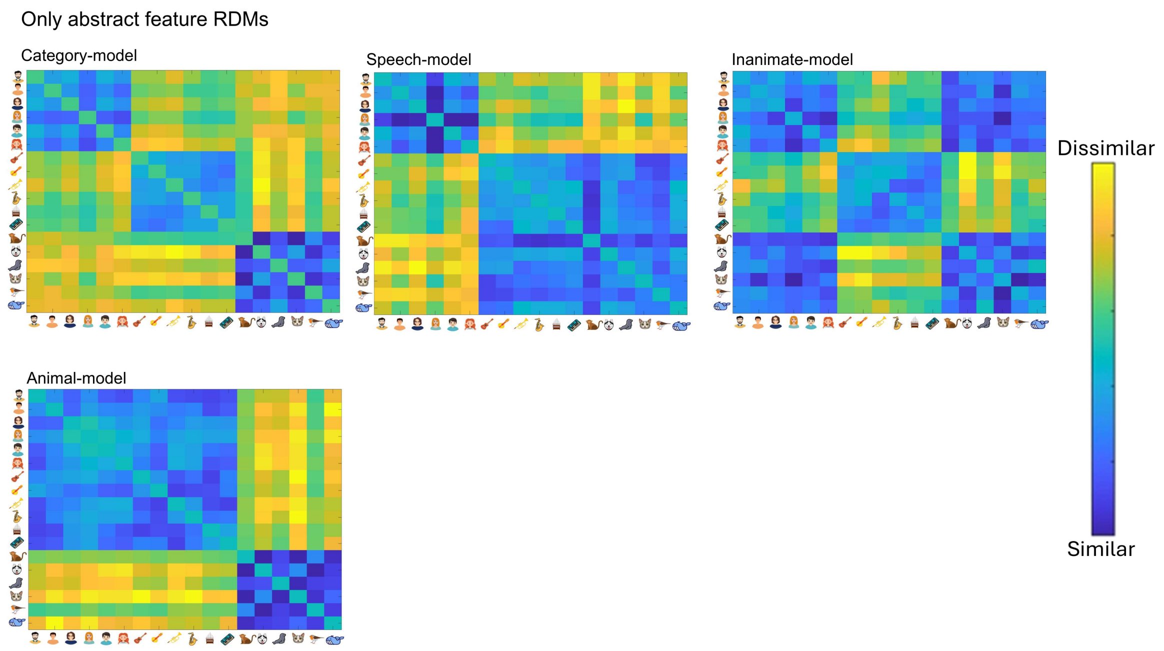


**Figure S8**. *Only abstract sound feature representational dissimilarity Matrices*. Here displayed are all 4 *Only abstract* sound feature RDMs. These RDMs are independent of all low-level sound feature models used in this study. Yellow colours indicate high dissimilarity between two subcategories for the given feature, whereas blue colours indicate high similarity.

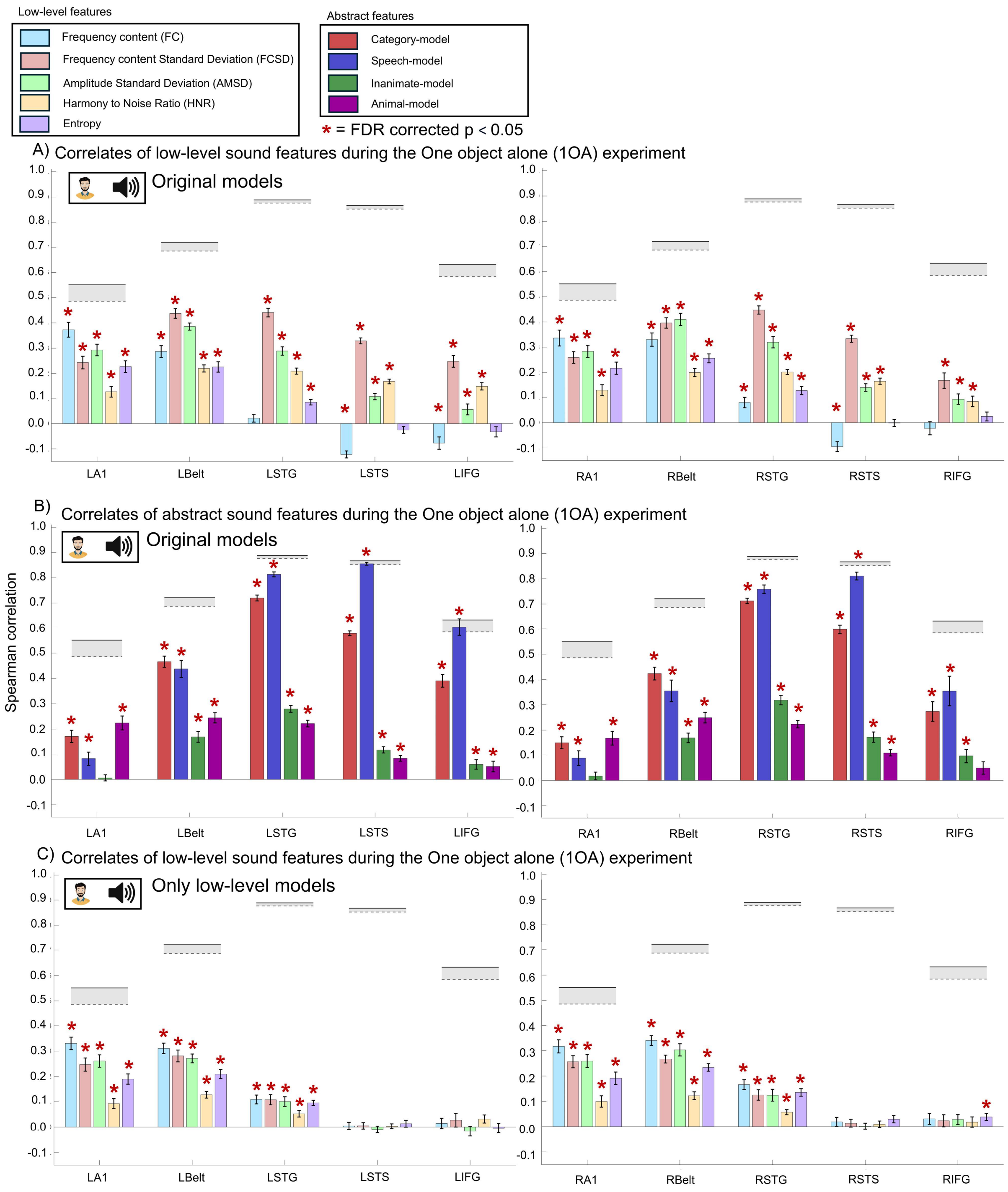


**Figure S9**. *ROI analysis for the one-object-alone experiment using Original and Only low-level models.* Bars show Spearman correlations between ROI representational dissimilarity matrices (RDMs) and sound-feature RDMs during the 1OA experiment. A) Displays correlations for low-level sound features using the *Original* models, which retain covariance with abstract sound-feature models. B) Shows correlations for abstract sound features using the *Original* models. C) Displays correlations for low-level sound features using the *Only low-level* models, in which variance shared with abstract models has been removed. Left and right columns show left- and right-hemisphere ROIs, respectively. Red asterisks indicate correlations that significantly deviate from zero (FDR-corrected p < 0.05). Error bars represent ±1 SEM. Grey bars denote estimated noise ceilings, with dashed and solid lines indicating lower and upper bounds, respectively.

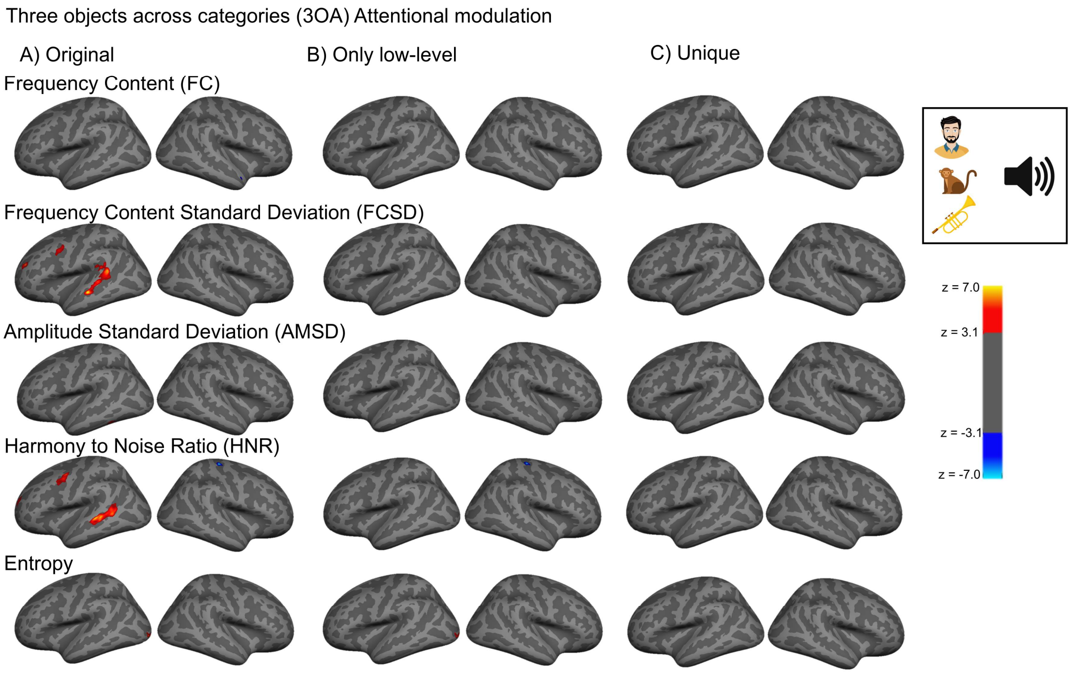


**Figure S10.** *Attentional modulation of low-level sound features during the 3OA experiment.* This figure shows attentional modulation of low-level sound-feature processing during the 3OA experiment. Warm colors indicate regions where the correlation between the attended-sound RDM and a given feature-RDM is significantly greater than the corresponding correlation for the distractor-sound RDM; cool colors indicate the opposite pattern. Significant attentional modulation was observed only when using the *Original* models. Because the *Original* models share covariance with abstract features, these effects are likely attributable to that shared variance, as no attentional modulation emerged for the residualised models.

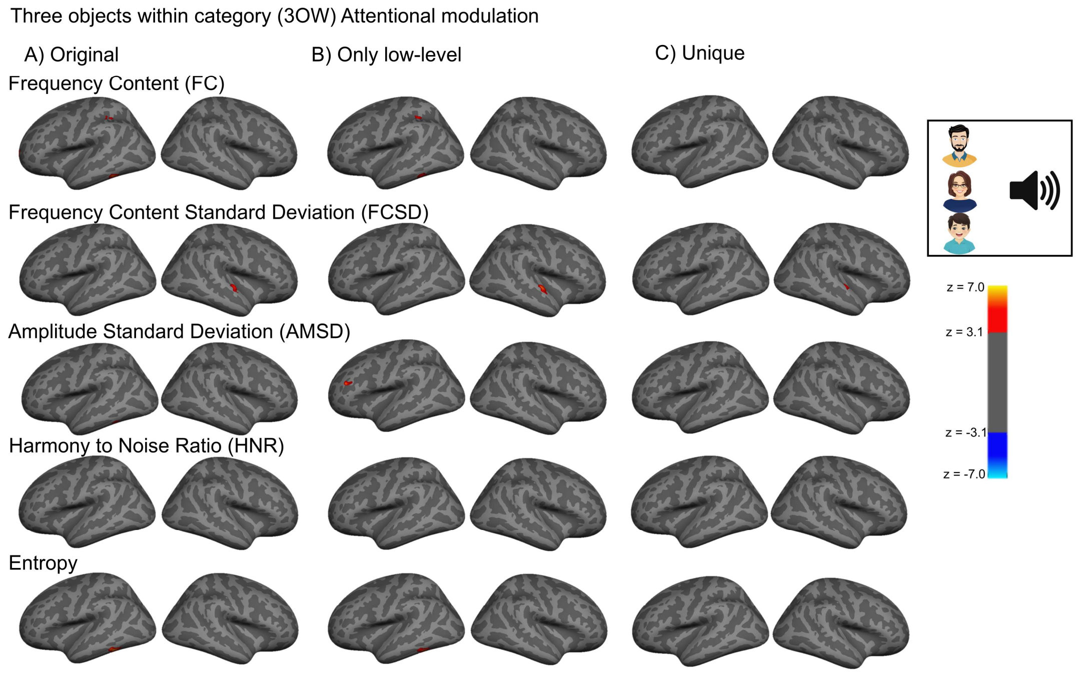


**Figure S11.** *Attentional modulation of low-level sound features during the 3OW experiment.* This figure shows attentional modulation of low-level sound-feature processing during the 3OW experiment. Warm colors indicate regions where the correlation between the attended-sound RDM and a given feature-RDM is significantly greater than the corresponding correlation for the distractor-sound RDM; cool colors indicate the opposite pattern.

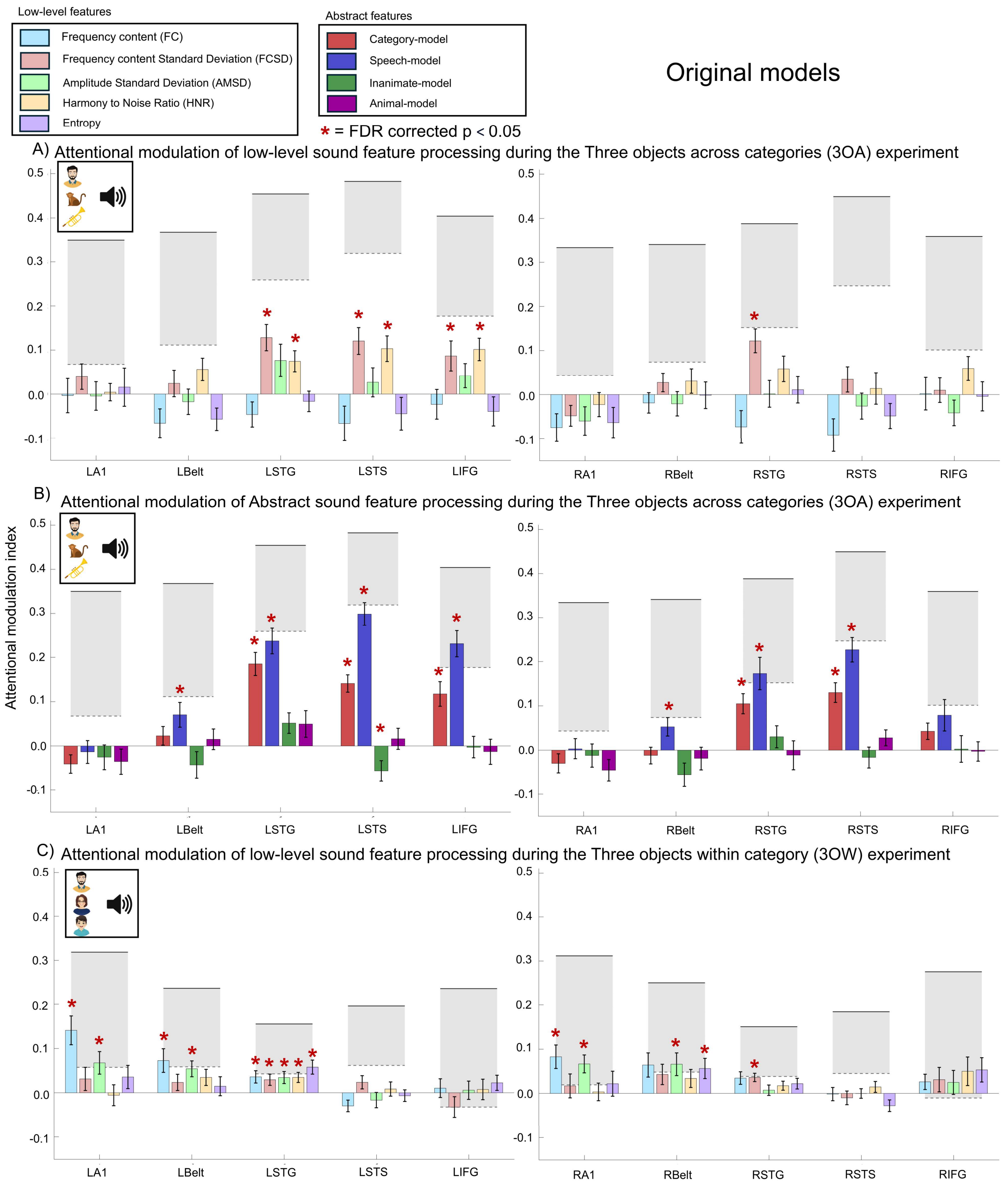


**Figure S12.** *ROI analysis of attentional modulation of sound feature processing during selective attention using Original models.* Attentional modulation indices (attended minus distractor correlations; see Figure 4) are shown for low-level and abstract sound features during the selective-attention experiments. A) Attentional modulation of low-level sound feature processing during the Three Objects Across Categories (3OA) experiment. B) Attentional modulation of abstract sound feature processing during the 3OA experiment. C) Attentional modulation of low-level sound feature processing during the Three Objects Within Category (3OW) experiment. Bars represent mean attentional modulation indices for each ROI, shown separately for left and right hemispheres. Red asterisks indicate values that significantly deviate from zero (FDR-corrected $p < 0.05$). Error bars represent ±1 SEM. Grey bars denote estimated noise ceilings, with dashed and solid lines indicating lower and upper bounds, respectively.

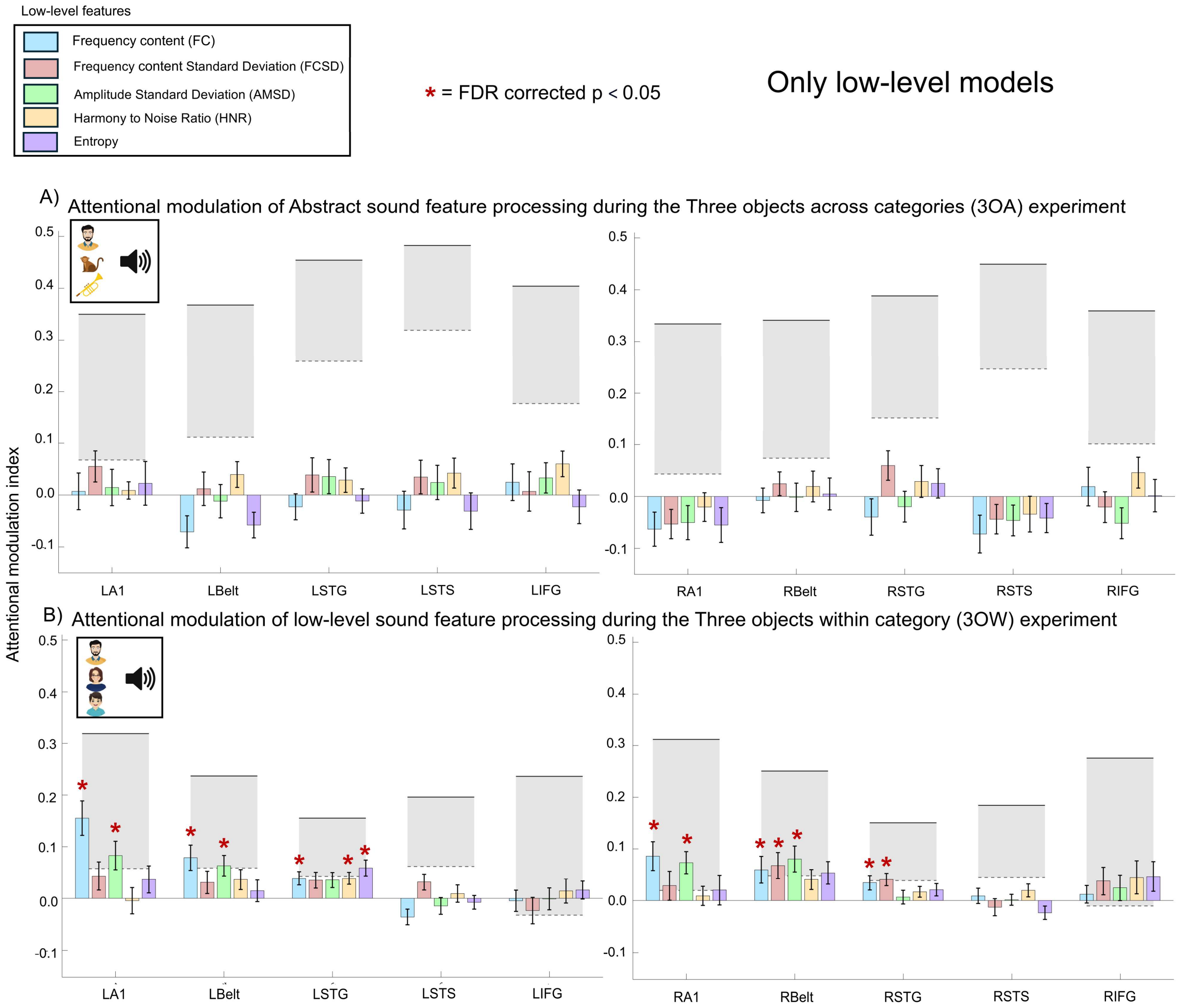


**Figure S13.** *ROI analysis of attentional modulation of low-level sound feature processing using Only low-level models.* Attentional modulation indices (attended minus distractor correlations; see Figure 4) are shown for low-level sound features during the selective-attention experiments using Only low-level models, in which variance shared with abstract sound-feature models has been removed. A) Attentional modulation of low-level sound feature processing during the Three Objects Across Categories (3OA) experiment. B) Attentional modulation of low-level sound feature processing during the Three Objects Within Category (3OW) experiment. Bars represent mean attentional modulation indices for each ROI, shown separately for left and right hemispheres. Red asterisks indicate values that significantly deviate from zero (FDR-corrected $p < 0.05$). Error bars represent ±1 SEM. Grey bars denote estimated noise ceilings, with dashed and solid lines indicating lower and upper bounds, respectively.
